# Optimising machine learning prediction of minimum inhibitory concentrations in *Klebsiella pneumoniae*

**DOI:** 10.1101/2023.11.20.567835

**Authors:** Gherard Batisti Biffignandi, Leonid Chindelevitch, Marta Corbella, Edward Feil, Davide Sassera, John A. Lees

## Abstract

Minimum Inhibitory Concentrations (MICs) are the gold standard for quantitatively measuring antibiotic resistance. However, lab-based MIC determination can be time-consuming and suffers from low reproducibility, and interpretation as sensitive or resistant relies on guidelines which change over time.

Genome sequencing and machine learning promise to allow in-silico MIC prediction as an alternative approach which overcomes some of these difficulties, albeit the interpretation of MIC is still needed. Nevertheless, precisely how we should handle MIC data when dealing with predictive models remains unclear, since they are measured semi-quantitatively, with varying resolution, and are typically also left- and right-censored within varying ranges.

We therefore investigated genome-based prediction of MICs in the pathogen *Klebsiella pneumoniae* using 4367 genomes with both simulated semi-quantitative traits and real MICs. As we were focused on clinical interpretation, we used interpretable rather than black-box machine learning models, namely, Elastic Net, Random Forests, and linear mixed models.

Simulated traits were generated accounting for oligogenic, polygenic, and homoplastic genetic effects with different levels of heritability. Then we assessed how model prediction accuracy was affected when MICs were framed as regression and classification.

Our results showed that treating the MICs differently depending on the number of concentration levels of antibiotic available was the most promising learning strategy.

Specifically, to optimise both prediction accuracy and inference of the correct causal variants, we recommend considering the MICs as continuous and framing the learning problem as a regression when the number of observed antibiotic concentration levels is large, whereas with a smaller number of concentration levels they should be treated as a categorical variable and the learning problem should be framed as a classification.

Our findings also underline how predictive models can be improved when prior biological knowledge is taken into account, due to the varying genetic architecture of each antibiotic resistance trait. Finally, we emphasise that incrementing the population database is pivotal for the future clinical implementation of these models to support routine machine-learning based diagnostics.

**Data Summary:** The scripts used to run and fit the models can be found at https://github.com/gbatbiff/Kpneu_MIC_prediction. The Illumina sequences from Thorpe et al. are available from the European Nucleotide Archive under accession <u>PRJEB27342</u>. All the other genomes are available on https://www.bv-brc.org/ database.

**Impact statement:** *Klebsiella pneumoniae* is a leading cause of hospital and community acquired infections worldwide, highly contributing to the global burden of antimicrobial resistance (AMR).

Ordinary methods to assess antibiotic resistance are not always satisfactory, and may not be effective in terms of costs and delays, so robust methods able to accurately predict AMR are increasingly needed. Genome-based prediction of minimum inhibitory concentrations (MICs) through machine learning methods is a promising tool to assist clinical diagnosis, also offsetting phenotypic MIC discordance between the different culture-based assays.

However, benchmarking predictive models against phenotypic data is problematic due to inconsistencies in the way these data are generated and how they should be handled remains unclear.

In this work, we focused on genome-based prediction of MIC and evaluated the performance of interpretable machine learning models across different genetic architectures and data encodings. Our workflow highlighted how MICs need to be treated as different types of data depending on the method used to measure them, in particular considering each antibiotic separately. Our findings shed further light on the factors affecting model performance, paving the way to future improvements of antibiotic resistance prediction.

## Introduction

Antimicrobial resistance (AMR), a major threat to human health worldwide, estimating around one million deaths are directly attributable to bacterial AMR every year [1]. AMR can be caused by single genes, or multiple loci can be involved [2]. For example, carbapenem non-susceptibility can be conferred by the production of a single carbapenemase enzyme encoded by single genes (eg *bla*_KPC_). However, synergistic effects such as the inactivation of an outer membrane protein or overexpression of efflux pumps can occur, in resistance associated to polygenic traits [3], where effect sizes greatly vary across the involved genes. Another example of synergistic effects is the epistatic interaction between *pbp* loci encoding for the penicillin-binding proteins (*pbp2x*, *pbp1a*, and *pbp2b*) modulates beta-lactam resistance in pneumococci [4, 5].

Indeed, antibiotic resistance is not strictly binary – different degrees of antibiotic resistance exist. Microbiologists measure the degree of resistance using the Minimum Inhibitory Concentration (MIC), a measure of the concentration at which the antibiotic inhibits bacterial growth in standard culture conditions. Several MIC measuring methods exist, differing by the antibiotics and range of concentrations tested. For example, solid-based methods have an extended range of MICs compared to broth dilution, but are long and costly, and are limited to testing for one antibiotic at a time. MIC interpretation is based on threshold values called breakpoints, typically decided for each pathogen-antibiotic combination by experts at international organisations such as EUCAST (http://www.eucast.org) in Europe or CLSI (https://clsi.org/) in North America. This interpretation changes over time due to the guideline updates, making some time-series analyses inconsistent.

Whole genome sequencing (WGS) is a mature technology that can quickly and reliably provide information about complete genomes, allowing to build models to predict resistance based on the genomic variants [6, 7]. WGS can also resolve phenotypic discordance between different MIC assays [8–11] even showing predicted MIC as more reliable in some contexts [12–14]. WGS-based AMR prediction typically requires specific manually-curated catalogues of mutations produced by carefully designed bottom-up literature searches [15–17]. However, Genome Wide Association Studies (GWAS) have also been used for producing such catalogues [17, 18].

The application of supervised-machine learning has been introduced in genome-based diagnostics to build more accurate, although potentially less interpretable, predictors of antimicrobial susceptibility [19, 20]. However, model interpretability is key in the clinical context where the detection of specific genetic determinants of resistance is expected in order to provide confidence in the results’ accuracy. Therefore, the use of black-box algorithms has received only limited practical interest in clinical settings. Indeed, prediction of resistance has exploited several interpretable machine learning models such as Gradient Boosting [19, 21], Random Forests [22] and regularised linear regression [14]. These models already achieved promising results in the prediction of quantitative traits in several bacterial species such as *M. tuberculosis* (>93% accuracy for first-line drugs) [23], *K. pneumoniae* (92% accuracy)[24], and nontyphoidal *Salmonella* (95% accuracy) [21]. Somewhat less accurate predictive models have also been built for *A. baumannii*, *S. aureus*, *S. pneumoniae* (accuracy range 88-99%) [19, 22].

However, benchmarking these models is challenging since their accuracy can be affected by population structure and the different genetic architecture of each resistance trait [25], potentially leading to false positive associations in highly clonal populations [26–29]. Another limitation in these studies is caused by the specifics of MIC measurement and interpretation, which may affect the prediction accuracy and genotype-phenotype correlation [30]. Due to the semi-quantitative nature of MICs and the limits on varying antibiotic-step concentrations, they can be considered censored, specifically right-censored (MICs greater than the last concentration tested) and left-censored (growth inhibition at the lowest concentration tested). Furthermore, even within the range tested, the actual MIC value is not known precisely, as it lies between two consecutively tested concentrations (dilutions). Therefore, how to handle MIC data for training models has not been fully addressed, limiting predictive model accuracy.

The aim of our work is to investigate the effect of MIC encoding (categorical or numerical, censored or not) on machine learning’s ability to predict them from WGS data. More specifically, we investigated the effects of the number of dilutions tested and the encoding of MICs as numeric or categorical/ordinal data on the prediction. We used *Klebsiella pneumoniae* as a case study, and tested the accuracy of machine learning predictions in several simulated phenotypic scenarios and on a real MIC dataset, highlighting the importance of MICs measurement representation in such analyses.

## Methods

### A combined dataset of MICs in *Klebsiella pneumoniae*

To ensure a realistic and representative basis for the quantitative trait simulation, a total of 4367 *K. pneumoniae* genomes were collated using three publicly available datasets [24, 31, 32]. These datasets include MIC phenotypes, with measurements for four antibiotics on the strains from Thorpe et al reported here for the first time. The MICs for all three collections have been determined by broth microdilution; those for the Thorpe et al. and Nguyen et al. collections, using the BD-Phoenix system (BD Diagnostics, Sparks, MD, USA), and those for David et al, using lyophilized custom plates (Thermofisher Scientific). The full dataset includes isolates sampled between 2011 and 2018 worldwide from human, livestock, and environmental samples, obtaining a collection representative of *K. pneumoniae* diversity.

The core and accessory distances between bacterial genomes were calculated using the PopPUNK v2.4.0 [33] software, choosing the DBSCAN clustering option to fit the model. The 347 clusters defined by PopPUNK v2.4.0 were used to adjust for population structure in the downstream analyses. Two major clusters, PP1 and PP2, contained about 30% of the dataset.

### Genotype data for quantitative phenotype simulation

To preserve the complex genetic architecture of the *K. pneumoniae* species, we simulated quantitative traits based on real observed genotypes. To generate the data for the gene-based trait simulation, gene sequences were annotated using Prokka v1.14.6 [34] and then clustered with Panaroo v1.2.8 [35] in moderate mode, giving a pan-genome with a total of 35380 genes. The presence absence matrix obtained from Panaroo v1.2.8 was converted to a VCF format by selecting the loci with minor allele frequency (MAF) > 0.5% (11961 genes) using PLINK v1.90 [36].

SNP calling was performed as follows: the core SNPs were called using Purple v1.22.2 [37] using the *K. pneumoniae* strain 30660/NJST258_1 as reference, and an annotated VCF file was generated.

To generate the input data for the GWAS simulation using SNPs, only biallelic (core-SNPs) in the VCF file with MAF>0.5% and Linkage Disequilibrium (LD) < 0.6 were selected as input for the phenotype simulation using BCFtools (6295 SNPs). Since linkage disequilibrium can span long genomic distances in bacteria [29], the LD window was calculated on the entire genome (~5Mb), as recommended for highly-recombinant species [38]. This filtering step prevents strongly associated loci from being included in the phenotype simulation as causal markers.

### Simulating phenotype data from real genotypes

To evaluate the predictive performance of each statistical model, quantitative MIC phenotypes were generated from the real genotypes in four different ways. Both genes and SNPs were used in the simulations. Simulated phenotypes were generated via the Genome-wide Complex Trait Analysis (GCTA) v1.93.3 [39]. Separate simulations for oligogenic, polygenic and homoplastic traits were carried out, as detailed below. Four different simulations were performed, using different values for the effect sizes (ES) — the contribution of a specific locus to the genetic variance of the trait - and narrow-sense heritability (*h^2^*) - the total proportion of variance of a trait explained by additive genetic effects.

To observe how the simulated phenotypes were affected by the presence/absence of the trait over different *h^2^* and ES levels, two preliminary simulations were performed using two known causal markers of beta-lactam resistance (KPC and CTX-M). The presence/absence of beta-lactamase genes predicted by Panaroo were manually compared with Kleborate v2.1 [40] and corrected if necessary. The obtained quantitative phenotypes were then rescaled to the [−1,1] interval.

After the preliminary simulations, a total of four different GWAS simulations were performed.

1. An oligogenic simulation selected three truly causal genes each with equal effect sizes 1.5, 2.5, 10, 30 and 100 [26, 41, 42]. Three genetic markers *bla*CTX-M, *bla*OXA and *bla*KPC involved in beta-lactam resistance were chosen to be causative of the trait.
2. A polygenic simulation selected 1000 random SNPs across the genome with same [43];
3. A homoplastic simulation [44] selected one causal SNP exhibiting homoplasy. Homoplastic sites were detected with HomoplasyFinder v0.9 [45]. The SNP-based phylogeny required as input to HomoplasyFinder was inferred with IQ-tree v2.2.0 [46], choosing the General Time Reversible (GTR) model with ascertainment bias with the Lewis [47] correction and 1000 ultrafast bootstraps. The causative homoplastic SNP within the *maoA* gene which encodes a positive regulator of the monoamine oxidase (see Supplementary Figure 1 for SNP distribution across the phylogeny), was chosen according to the consistency index (CI=0.023), indicating high homoplasy;
4. An oligogenic simulation [48] selected eight different SNPs with varying effect sizes (range −0.39:8.18) and homoplasy (CI range 0.008:0.66) between the two groups. The homoplastic sites were detected as described above in the homoplasy simulation.

### Converting simulated phenotypes to MIC values

The output of the phenotype simulations is a quantitative, continuous trait. However, in reality the MIC is measured semi-quantitatively, resulting in the measurement of MIC as an approximation of the exact inhibitory concentration of the antibiotic due to the limited precision of double dilutions. Furthermore, left and right censoring of the true MIC occurs at the ends of the testing range. The range of concentrations tested by the broth-based instrument typically includes between 3 and 7 doublings. We therefore empirically binned the simulated continuous traits into 4 or 6 equal intervals. In addition, we also resemble the E-test method to be used when testing the regression model binning the quantitative traits into 4, 6, 8 and 10 categories.

The censoring can improve the class balance e.g. by incorporating the minimum and maximum at the end of testing range values within the nearest bin (Supplementary Figure 2).

In addition, the midpoint of each bin (not converted to log) was used as the phenotype with both a classification and regression problem. In case of regression the bins were also censored, incorporating the values upstream and downstream using the quantile intervals (0.025,0.975). Lastly, when dealing with the real MICs, the >, <, ≥, and ≤ symbols were removed and the values converted to log_2_ values, as previously shown [24, 30]. The log_2_ values of the filtered MICs were used as labels for all machine learning tasks.

### Machine learning models

We used two machine learning models: Elastic Net and Random Forests. These were chosen due to their interpretability and scalability as well as accuracy [[28, 49], 20]. In addition, both these models allow us to apply population structure correction, as explained below.

To evaluate the performance of statistical models in detecting the causal markers, Elastic Net and Random Forest were tested on the simulated data, using independently two presence-absence matrices as input, coreSNPs- and genes-based respectively. Both regression and classification setups were used, allowing us to highlight how the framing of statistical problems affected the predictive accuracy of the models.

All the analyses run using Elastic Net and Random Forest were performed by splitting the dataset randomly into 70% training and 30% testing.

Regression and multinomial classification performance was assessed using the proportion of variance explained (R^2^) and the balanced accuracy (bACC), respectively. Classification accuracy was also evaluated using a ± 1 two-fold dilution factor [30] a more flexible measure of accuracy especially when there are many ordered classes (concentrations).

#### Elastic Net

Elastic Net [50] is a penalized linear regression model that includes both the L1 (LASSO) and L2 (Ridge) penalties to the loss function during training. The L1 penalty shrinks the coefficients towards zero, which removes many predictors from the model. The L2 penalty instead minimises the Euclidean norm of the coefficient vector, but typically produces models that use all the predictors. Elastic Net can successfully manage dependent variables, expected due to a strong linkage disequilibrium in bacterial populations [26]. The model was tested using α = 0.01 [28], λ = lambda.1se, indicating the largest value of *λ* whose cross-validated error falls within 1 standard error of the minimum such error. A 10-fold cross-validation was used on the training data. The model was run using the “glmnet” package [51] in R software v4.1.3.

#### Random Forest

Random Forest is a tree ensemble method that provides an improvement over bagged trees by decorrelating the trees, reducing the variance. We expected Random Forest to successfully handle both multiclass problems and multicollinearity [52]. The Random Forest model was trained using 500 trees, and the Gini index for classification and the variance of the responses for regression as the impurity measure, respectively. The Random Forest model was run using the “ranger” library [53].

### Computation time

We also compared the performances of the two models in terms of computational resources (Supplementary Figure 16) using 30 threads on a server with 504Gb memory and an Intel Xeon CPU E5-2699 v4 2.20GHz.

### GWAS

GWAS is a tool frequently used to unveil the genetic causes associated with a phenotype of interest, as well as to study the heritability of complex traits such as antibiotic resistance.

Whereas using machine learning approaches use all genetic variants to fit a model predicting the phenotype, GWAS analysis infers individual associations between genotype and phenotype. To turn GWAS results into a phenotype predictor, we used the standard approach of polygenic risk scores. We selected all the variants with a statistically significant association, and obtained the predictor by weighting them by the coefficients obtained from the regression.

GWAS was run using the Pyseer software [28], selecting the Linear Mixed Model of fixed and random effects. This model is based on the FaST-LMM’s [54] likelihood calculation in linear time for each variant, therefore the variant effects are calculated as marginal effects, rather than in a joint model as for the above two methods.

Also, we assessed a prediction with FaST-LMM exploiting the GWAS results by selecting the significant variants at threshold α=0.05 after adjusting for multiple testing using Bonferroni correction (adjusted p-value threshold = 5.5E-06), using as number of tests the amount of SNPs detected. Although Bonferroni is a highly conservative method that potentially suffers from false negative rate when variants are not independent, it works properly when the selected loci are not in strong linkage disequilibrium i.e. LD<0.6.

### Population structure correction

Due to the complex dynamics of bacterial populations (e.g. lineage effects), the models were tested adjusting for population structure. Sequence reweighting [28] was used when running the Elastic Net and Random Forest with the clusters assessed by PopPUNK to incorporate the effects of bacterial strains. This gives a weight inversely proportional to the cluster size to each observation within a cluster. The reweighting therefore allows the model to account for all observations by giving less importance to those belonging to the same lineages.

The adjustment for population structure in the FastLMM used for GWAS was performed through the kinship matrix (calculated from SNPs). This matrix accounts for genetic relatedness between strains and is included in the regression as a random effect.

## Results

### Simulating semi-quantitative traits with varying genetic architecture

In order to generate multiple realistic quantitative traits which account for different genetic scenarios, a total of four simulations were carried out, accounting for oligogenic, homoplastic and polygenic effects (See methods).

To observe the distributions of the simulated phenotype across different levels of *h*^2^ and effect sizes, an initial phenotype simulation was performed (Figure 1; Supplementary Figure 2) using two genes with fixed effect sizes (effect size = 2.5), as in the oligogenic simulation. The genes CTX-M-15 and KPC were selected as the causal variants according to clinical relevance due to their involvement in resistance to beta-lactams and carbapenems.

**Figure 1).**
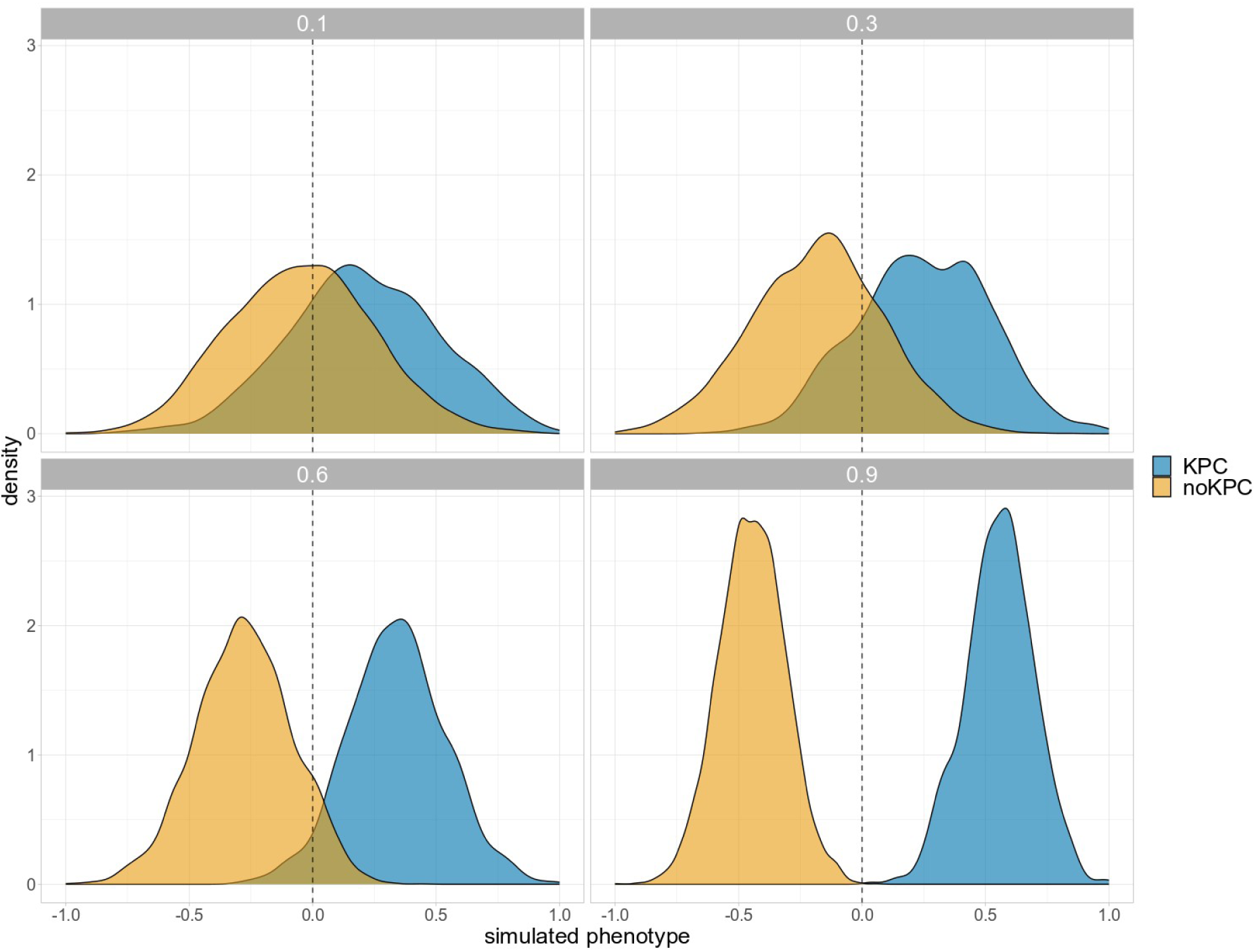
Simulated phenotype distribution using (a) non-homoplastic blaKPC gene as causal variants at different levels of h^2^. The areas are coloured according to the presence-absence of the causal variant.

Heritability related to AMR traits has been estimated to range between 0.4 to 0.9 depending on the genetic background of organisms, resistance traits and the model chosen [55]. Due to lack of knowledge about heritability in *K. pneumoniae* associated to AMR, we relied on previous studies on other organisms like *N. gonorrhoeae* and *S. pneumoniae* whether heritabilities of the different antibiotics resistance were high (e.g. *h^2^* > 0.6), both using binary or continuous phenotypes [56]. We noted that different effect sizes used in this simulation do not significantly affect the density distributions of the simulated phenotypes, with heritability having a more important effect over this range. Since this preliminary analysis on our dataset confirmed that the separation of the trait by marker started at *h^2^*=~0.6, the quantitative traits through the four simulations were generated starting from high levels of *h^2^*.

### Benchmark models using simulated MICs data

Most machine learning methods rely on the assumption that observations and predictors are identically and independently distributed, which is rarely the case with genomic data, particularly highly structured bacterial populations. Therefore, accounting for population structure when using these models is recommended to avoid false positive and spurious associations [26].

In this work, we benchmarked Elastic Net, Random Forest, interpretable and flexible machine learning methods able to handle high dimensional data where *p* >> *N*, both with the capability for regression and classification. In addition, Fast-LMM (through Pyseer) was used for regression analysis only, since it does not deal with multinomial classification.

The simulated quantitative traits were binned into classes of equal length using either two — to simulate both the binary (resistant and sensitive) interpretation of the MIC — four and six bins, resembling a realistic partition of MIC classes in reality. Also, the performances of the models were evaluated on both uncensored and censored data from the simulations. The censoring was applied because when measuring MIC by serial dilution, we have start and end dilutions, without testing concentrations at levels beyond these limits.

We first tested the models on multiple bins to assess the effect of MIC measurement resolution, highlighting whether treating the simulated traits as categorical variables can lead to gains in prediction accuracy (Figure 2; Supplementary Figure 4-11). Our analysis showed a decrease of the accuracy, with further decreases with more bins. For this reason, we also assessed the off-by-one prediction accuracy — discussed below — allowing us to improve the models interpretability whether the range of antibiotic-step concentrations is broader.

**Figure 2).**
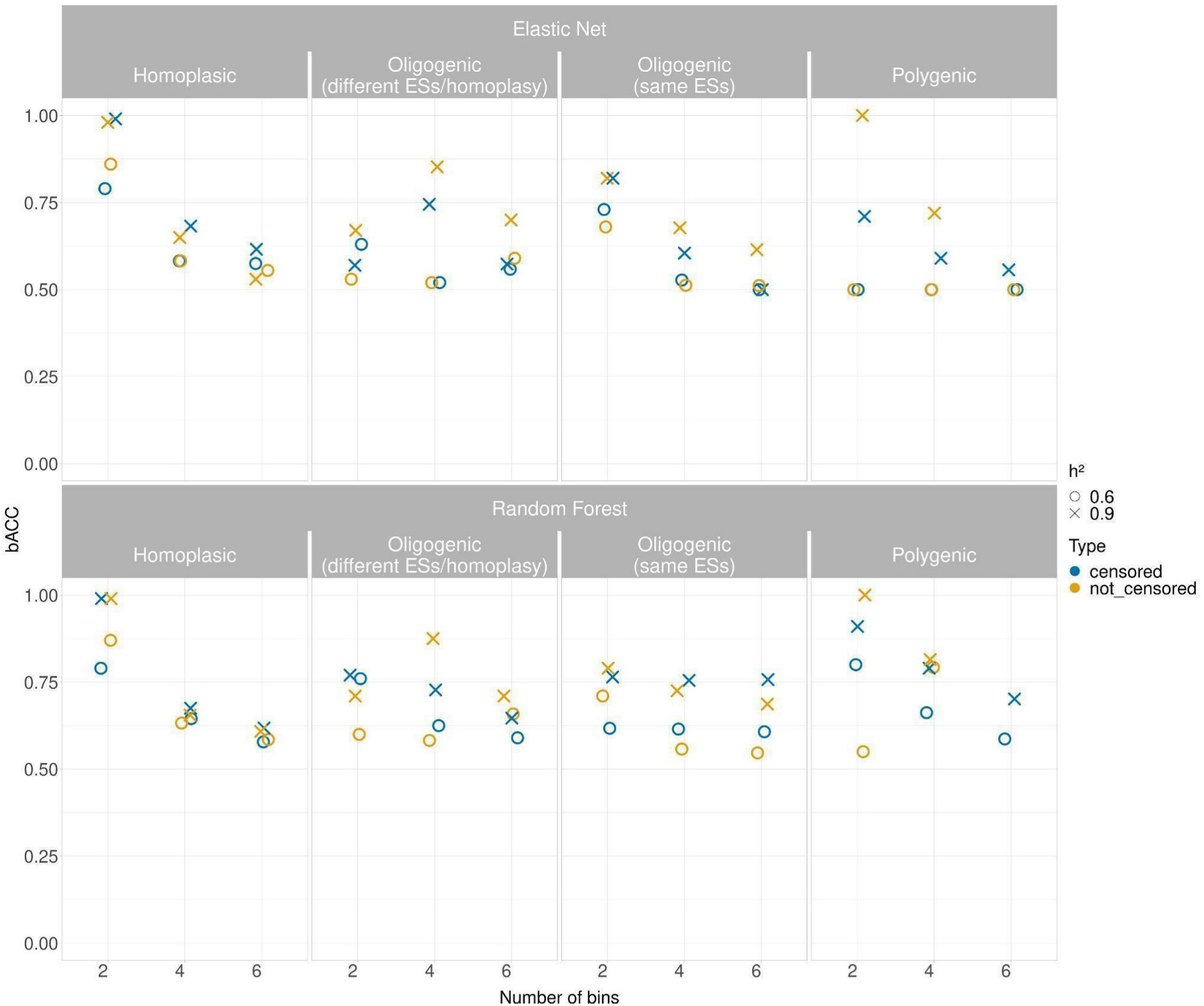
Performance of the classification models (effect size = 2.5 except for the oligogenic simulation) measured using balanced accuracy (bACC), indicating the arithmetic mean of sensitivity and specificity. The models were benchmarked over two levels of h^2^, considering both the censored and not censored binned simulated traits.

Specifically, when the binned quantitative simulated traits without applying the censoring were used (Figure 2), the Elastic Net and Random Forest perform similarly in the homoplasic simulation, where bACC range of Elastic Net was 0.55-0.88 (h^2^=0.6); 0.53-0.98 (h^2^=0.9) and the bACC range of Random Forest was 0.58-0.97 (h^2^=0.6); 0.60-0.99 (h^2^=0.9). A similar result was observed into the two oligogenic simulations where the Random Forest average bACC was slightly better in all the settings (range 0.68-0.87) compared to the Elastic Net (range 0.61-0.85) at h^2^=0.9. The prediction accuracy in polygenic simulation highlighted how Random Forest better handled the number of classes across the simulations (average bACC = 0.78) compared to the Elastic Net (average bACC = 0.68).

Since the true minimum inhibitory concentration may occur at the ends of the testing range in reality, we also assessed how censoring the simulated traits affected the accuracy of the models. The application of censoring can group classes that contain fewer observations by their incorporation within the nearest bin (Supplementary Figure 3) increasing class balance. However, this method may reduce the amount of information available to the model.

Indeed, in this section we observe the capability to assess the prediction in the polygenic simulation with six bins (*h^2^*=0.9) for both Elastic Net and Random Forest, not applicable in the first comparison without the censoring (Figure 2). Although we observed a increasing in accuracy only at *h^2^*=0.6, especially in case of two bins in the oligogenic (different effect size / homoplasy) (bACC = 0.76) and polygenic (bACC = 0.8), the accuracy was overall low when the number of the classes increased.

Since the accuracy was affected as the number of the bins increased, we also used a more flexible measurement of accuracy measuring how prediction deviates from the correct class, specifically allowing one class higher or lower as correct, as was previously suggested [30]. This measure allowed us to assess a more interpretable measure of accuracy, particularly when the number of classes increased.

In this setting (Figure 3; Supplementary Figure 8-11), we observed that the Random Forest achieved a higher accuracy in all simulations (range bACC = 0.86-1) regardless of the *h^2^* levels and in the presence of censoring (Figure 3). Albeit also the Elastic Net gained an improvement of the prediction accuracy where the traits were not censored (bACC range 0.88-0.99), its accuracy was consistently lower when the censoring was applied (bACC range 0.5-1).

**Figure 3).**
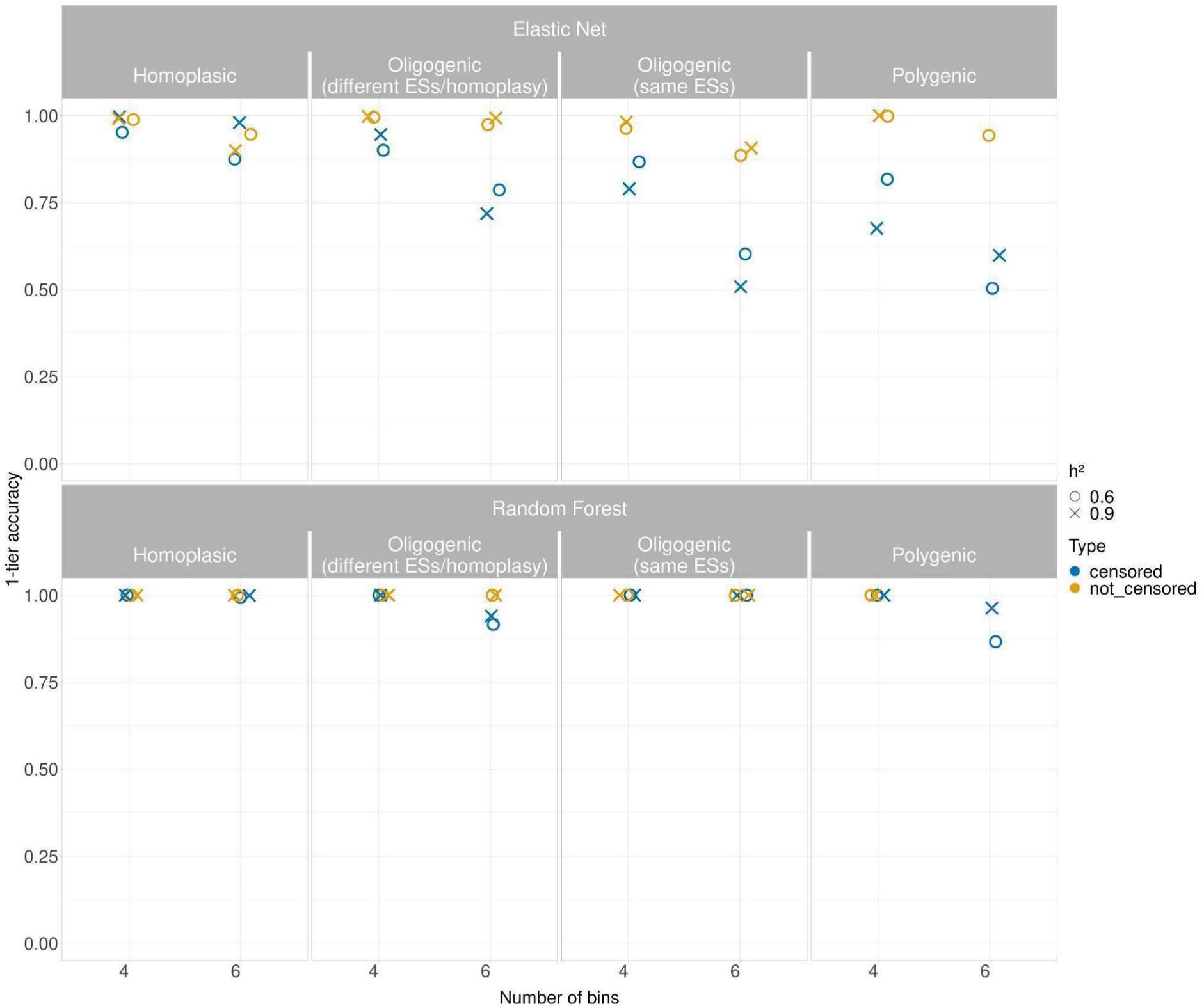
Performance of the classification models (effect size = 2.5 except for the oligogenic simulation) when the accuracy (need to correct y-axis label) was adjusted to consider one-either-side predicted class as correct (accuracy within ± 1 two-fold dilution factor). The models were benchmarked over two levels of h^2^, considering both the censored and not censored binned simulated traits.

Dealing with ordinal/categorical data usually adversely affects the accuracy of predictive models as the number of classes increase, as more misclassification categories become possible. As a large number of bins approaches a continuously distributed variable, we also assess how the performance of the models were affected when treating the simulated MICs as real numbers rather than categories (Figure 4; Supplementary Figure 12-15), as previous studies have already assessed [21, 24, 30].

**Figure 4).**
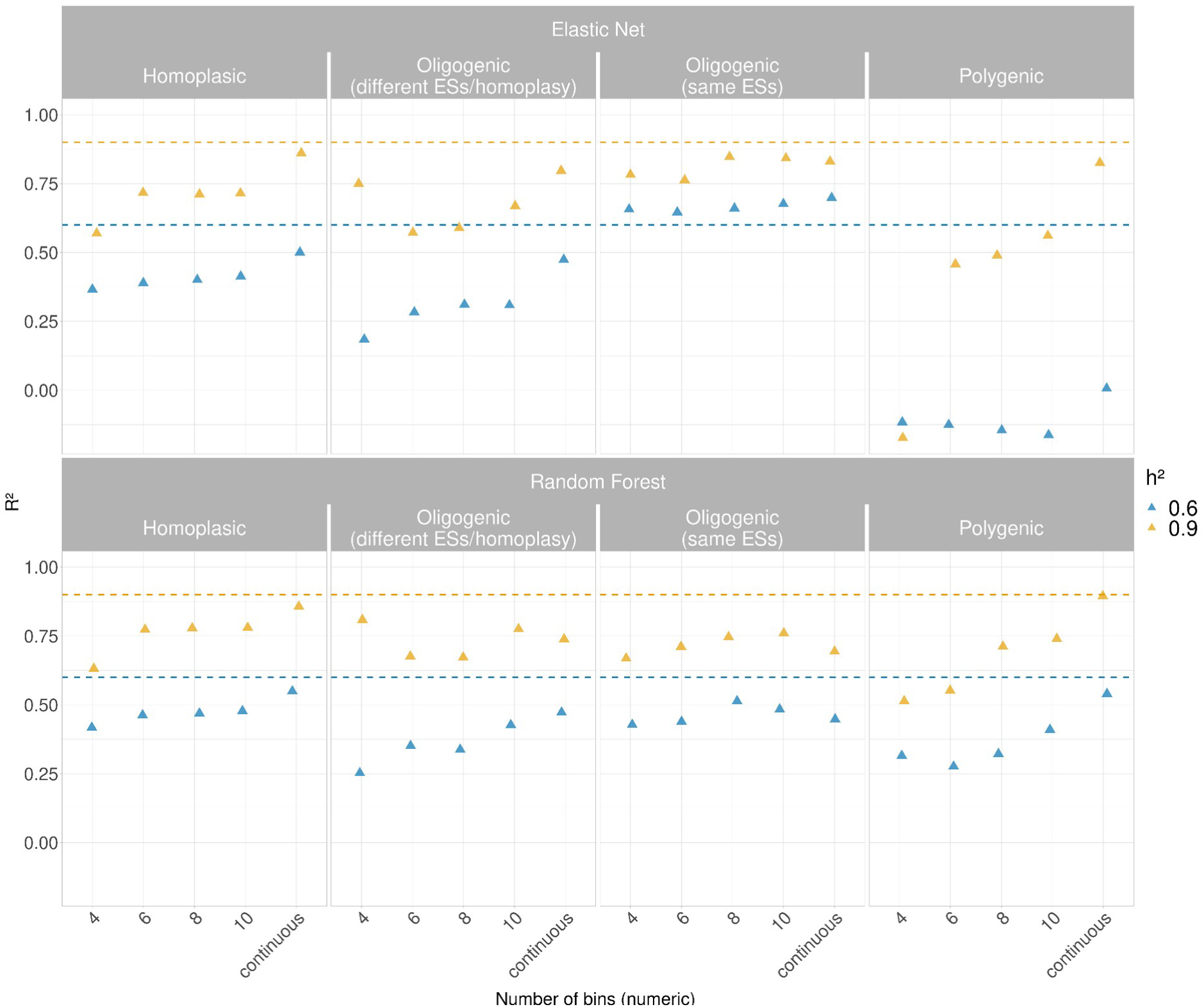
Performance of regression model (effect size = 2.5 except for the oligogenic simulation) between Elastic Net and Random Forest when dealing with simulated MICs. The bins obtained from the simulation were binned into multiple intervals and treated as numeric. Also, the simulated quantitative traits without binning were used. Since the h^2^ is the proportion of phenotypic variance explained by genotype, and thus an equivalent of the R^2^ in regression, the dashed lines are used to assess the capability of the models to estimate the h^2^.

In this setting, we considered 4, 6, 8 and 10 binned phenotypes of the simulations (see above) using the midpoint of each bin as a continuous trait instead of as categories (Figure 4). Here, we calculated the performance using the *R^2^* value. Simultaneously we estimated the *h^2^*, since it represents the proportion of variance that can be attributed to the variation of genetic effects and thus the equivalent to the *R^2^* in regression analysis.

Figure 4 shows the performance of the two methods over the two levels of *h^2^*, where the dashed lines indicated the *h^2^* levels used to generate the quantitative simulated MICs. We pinpointed that the *R^2^* achieved by both the models increased overall proportionally with the number of the bins up to continuous distribution, as expected.

There was an overlapping trend between the models in the homoplasic and oligogenic (different effect size / homoplasy) simulations at both the levels of *h^2^*, and we observed only one exception where Elastic Net outperformed Random Forest in the oligogenic simulation with the same effect size (*R^2^* range 0.64-0.67; *h^2^*=0.6 and 0.76-0.84; *h^2^*=0.9). However, Elastic Net suffers more in cases of polygenic effects where the regression was performed on the bins, especially at *h^2^*=0.6.

About the *h^2^* estimation, an overlapping trend between the models was observed (Figure 4), albeit the Random Forests exhibited an overall better fit compared with Elastic Net except for the oligogenic simulation (genes with same effect sizes).

To summarise, the results on simulated data showed that for classification the overall performance on binary classification achieves better results, while the performance decreases as the number of classes increases, as expected due to the increased difficulty of exact classification. In addition, the accuracy was slightly better when the censoring was not applied to the simulated traits. The Random Forest resulted to be more reliable by handling predictions in the vast majority of the simulations, highlighting this algorithm as more suitable when tackling with class imbalance.

### Computational and memory requirements

The Random Forest model handled the classification better, showing no significant difference in time among the different scenarios (Supplementary Figure 16).

Both models were faster on regression than on classification, ranging between 5-10Gb of memory used.

### Selection of true causal variants in GWAS and machine learning models of MICs

We performed three GWAS using Elastic Net, Random Forests and FastLMM benchmarking them by the detection of true and false positives across the SNPs-based simulations.

When running Elastic Net and Random Forests we accounted for population structure using sequence reweighting whilst a kinship matrix calculated from SNPs was used for the FastLMM.

In addition, we retrieved the predictors using the beta (effect size) values from Fast-LMM GWAS output to assess a prediction of the simulated traits (as performed above with the other two models), since Pyseer does not implement prediction mode for the Fast-LMM mode.

The obtained prediction showed how in all the simulations the accuracy was lower compared to the previously tested model (*R^2^* range 0.27-0.43; *h^2^*=0.9 and 0.18-0.30; *h^2^*=0.6).

However, through the tuning of parameters such as the **α** in Elastic Net (e.g. L1 regularization), the variable importance in Random Forest and p-value in Pyseer, we assessed the overlap between the significant SNPs detected and the causal variants set in the simulations.

Supplementary Figures 17-19 show the performance of the Elastic Net, Random Forest and Pyseer according to the values used for **α**, importance and p-value respectively.

Considering the ability to detect the True Positive (TP) variants, all the models were able to correctly predict the causative SNP in the homoplasic simulation despite the level of *h^2^* and the different tuning of the parameters, underlining how a sparser model does not affect the power accuracy when dealing with monogenic traits. In addition, the Random Forest achieved the lower rate of FP even with low importance cut-off, with a maximum of 22 (*h^2^*=0.6) and 17 SNPs (*h^2^*=0.9).

In the Polygenic simulation, an overlapping trend was observed for Elastic Net and Pyseer, albeit the latter was more able to maintain a higher detection of causative SNPs regardless of the p-value cut-off, especially when the *h^2^*=0.9.

The detection rate of true positives in Random Forest was more affected when increasing the cut-off related to the importance of coefficients, though it allows more reliability to handle the False Positive (FP) compared to the other two models. Notably, when the Elastic Net was shifted from α=0 (equivalent to a Ridge regression) to α=0.07, the number of TP was drastically reduced, indicating how even introducing small sparsity in the model can shrink a large amount of putatively related loci [28].

FaST-LMM maintained a higher number of true positives detected despite the increasing *p*-value threshold in the polygenic simulation, highlighting how LMM models handle polygenic effects, as previously described [57]. However, the number of false positives was consistently high, possibly due to that kinship matrix obtained from SNPs can suffer from spurious associations in presence of high clonality compared to the use of a phylogenetic tree to compute relatedness between samples [26, 58].

Lastly, regarding the oligogenic traits where a few loci with different effect size and homoplasy were involved, Elastic Net accounted for more true positives avoiding spurious associations as the L1 penalty increased, suggesting Elastic Net as preferable in this scenario.

### Model performance and *h^2^* estimation on real MIC data

Hereafter, we applied our testing framework to real MICs data (Supplementary Table 1), since we cannot vary the genetic effects such as effect size distribution and homoplasy. Therefore, we assessed the model performances in terms of accuracy by cross validation.

Since the *K. pneumoniae* strains were not always tested for the same antibiotics across the three datasets, we selected three antibiotic classes of interest where the MICs were largely available (Figure 5). Thus, Fluoroquinolones, Aminoglycosides and Beta-lactams were considered and a representative drug for each class was selected by including Gentamicin (GEN), Ciprofloxacin (CPFX) Piperacillin/Tazobactam (TZP) and Meropenem (MEM). The number of the strains tested for the chosen antibiotics were the following: Meropenem (4220/4367), Gentamicin (3161/4367), Ciprofloxacin (3158/4367) and Piperacillin/Tazobactam (3276/4367). For each antibiotic we observed a different range of step-concentrations (range 4-10) that we treated both as continuous and unordered classifications (Figure 5).

**Figure 5).**
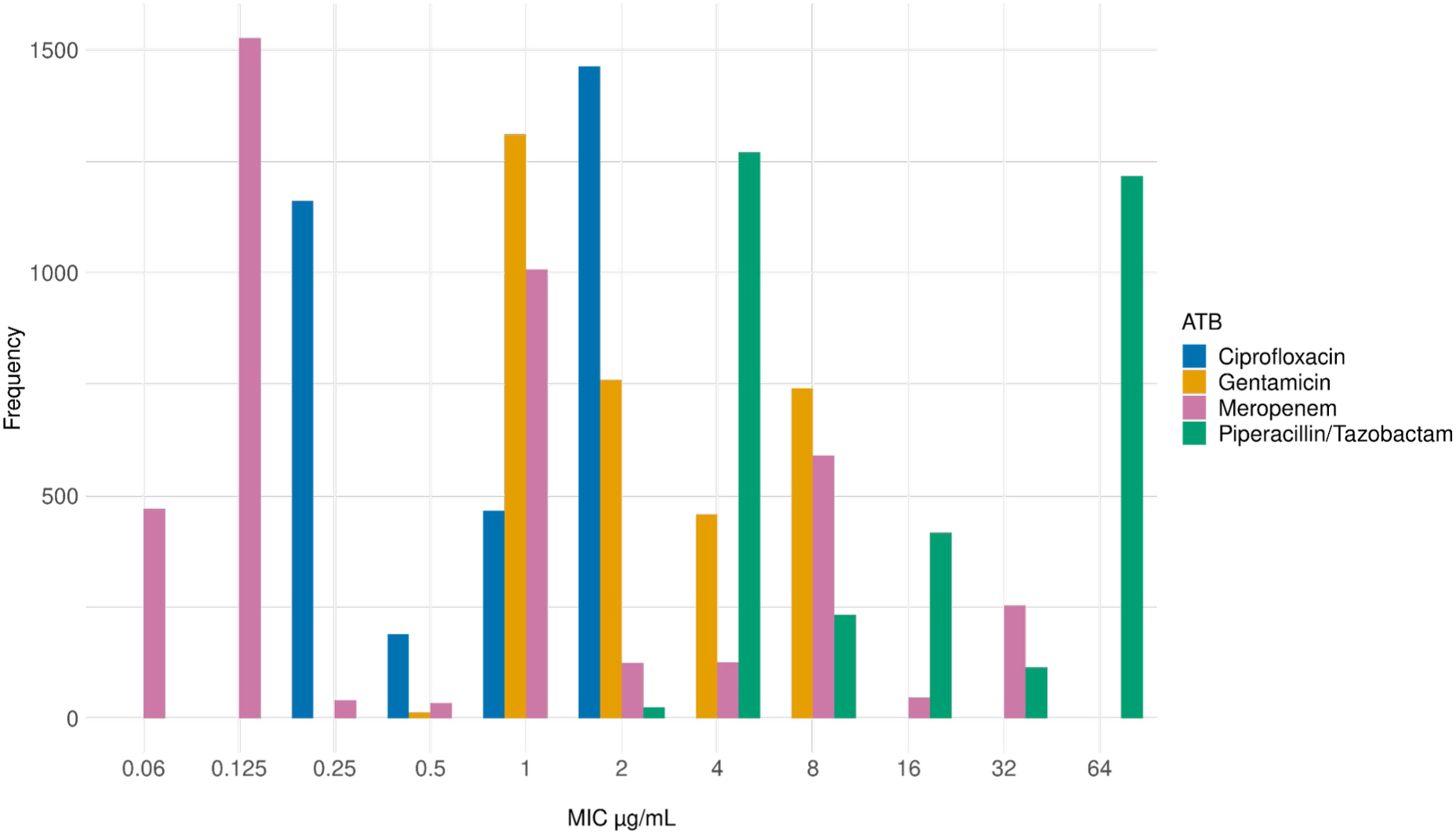
MIC values distribution among the different antibiotic classes

Dealing with MICs framed as classification (Table 1), Elastic Net and Random Forest showed comparable balanced accuracy when dealing with Gentamicin, Meropenem and Piperacillin/Tazobactam, while in case of Ciprofloxacin the accuracy was higher when using Elastic Net model. However, once the accuracy of classification prediction was measured by allowing one predicted class higher or lower as correct, we observed an increased performance in the accuracy of Elastic Net while the Random Forest did not exhibit any significant improvement, in contrast with what we observed in the simulations (Figure 3). When the MICs were framed as regression (Table 1), the Random Forest showed a better performance compared to the Elastic Net for all the antibiotics (*R^2^* range 0.51-0.72 vs 0.19-0.59), although it was overall low except in Ciprofloxacin (*R^2^*=0.72).

**Table 1).** Summary of the performances of the Elastic Net and Random Forest models using real MICs treated as unordered class and numerics. Four antibiotics (ATB), Ciprofloxacin (CPFX), Gentamicin (GEN), Piperacillin-Tazobactam (TZP) and Meropenem (MEM) were tested. Balanced accuracy (bACC) and the 1-tier accuracy (the inclusion of classes one-either-side as correct) were used for classification, and *R^2^* for regression, including also the one related to the train set. In addition, the computational performances in terms of time and the number of antibiotic concentration intervals were included.

| ATB | Model | bACC | ACC 1<br>error-side | Prediction<br>time (s) | Train/test<br>time<br>(s) | ATB<br>concentration<br>interval | R<sup>2</sup><br>train | R<sup>2</sup><br>test | Prediction<br>time (s) | Train/test<br>time (s) |
| --- | --- | --- | --- | --- | --- | --- | --- | --- | --- | --- |
| CPFX | Elastic<br>Net | 0.93 | 1 | 0.62 | 33.2 | 4 | 0.82 | 0.59 | 0.13 | 25.9 |
| CPFX | Random<br>Forest | 0.74 | 0.58 | 0.91 | 7.90 | 4 | 0.73 | 0.72 | 0.92 | 9.01 |
| GEN | Elastic<br>Net | 0.78 | 1 | 0.87 | 50.8 | 5 | 0.77 | 0.34 | 0.28 | 24.0 |
| GEN | Random<br>Forest | 0.69 | 0.40 | 0.85 | 7.87 | 5 | 0.53 | 0.51 | 0.89 | 8.73 |
| MEM | Elastic<br>Net | 0.70 | 0.89 | 3.46 | 2.97 | 10 | 0.51 | 0.21 | 0.39 | 31.4 |
| MEM | Random<br>Forest | 0.58 | 0.10 | 1.03 | 9.80 | 10 | 0.50 | 0.48 | 0.87 | 10.1 |
| TZP | Elastic<br>Net | 0.62 | 0.99 | 1.43 | 5.02 | 6 | 0.70 | 0.19 | 0.28 | 30.4 |
| TZP | Random<br>Forest | 0.64 | 0.99 | 0.62 | 7.47 | 6 | 0.59 | 0.57 | 0.93 | 8.61 |

In addition, both the Random Forest and Elastic Net showed better performance on the *R^2^* train set compared to the *R^2^* test set (Table 1), suggesting that these models are prone to overfitting.

The results on real MICs highlighted how the classification model outperforms the regression one, with the only exemption of Random Forest when the prediction of Ciprofloxacin was assessed, showing an overlapping accuracy between the two cases.

We also benchmark Elastic Net, Random Forest and Pyseer for the *h^2^* estimation (Figure 6). The *h^2^* associated with antibiotic resistance is expected to be high, on the basis that the trait is largely determined by highly penetrant additive genetic variants directly causal for the resistance mechanism [26]. Indeed, the *h^2^* estimation of the three models on Ciprofloxacin, Gentamicin and Piperacillin/Tazobactam indicates these traits as highly and moderately penetrant, whilst the estimated *h^2^* for Meropenem exhibits a lower level. These results suggested how the different intervals of estimated *h^2^* can be associated with the location of genetic causative variants of the antibiotic resistance. Indeed, when the resistance is mainly associated with genes carried on plasmids, the genetic variation can be poorly accounted (e.g. for in a kinship matrix), since information of causal variants is not always located in core positions.

**Figure 6).**
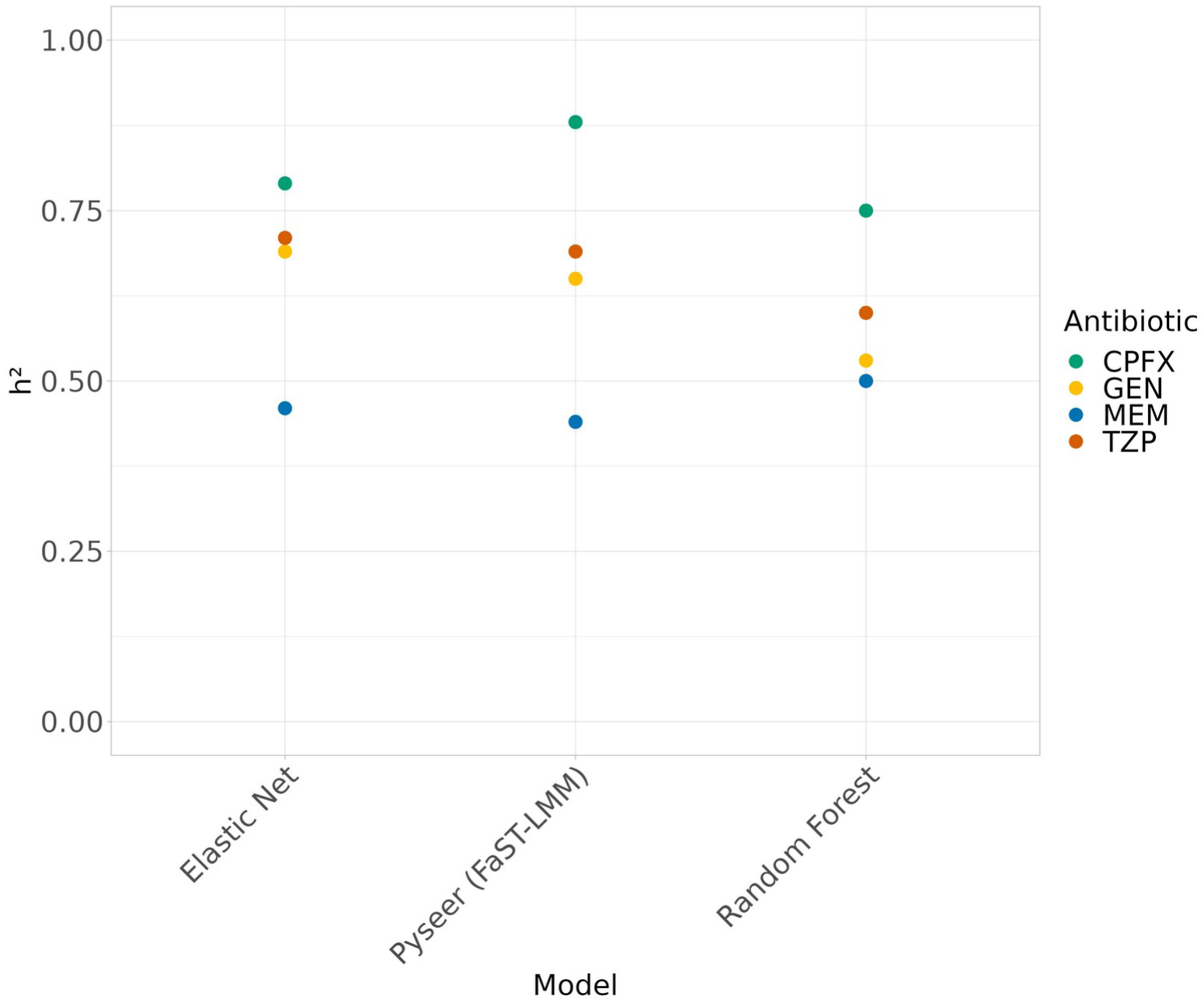
Heritability estimation of antibiotic resistances using real MICs. Four antibiotics Ciprofloxacin (CPFX), Gentamicin (GEN), Piperacillin/Tazobactam (TZP) and Meropenem (MEM) were tested.

## Discussion

Machine learning based prediction of MIC starting from genomic data offers an attractive supplement to the traditional phenotypic methods. However, how to specifically handle MICs data is still not addressed due to their semi-quantitatively measurement, including the left- and right-censoring within varying ranges.

In this work, we sought to investigate how the encoding of semi-quantitative resistance traits as MIC values affects the accuracy of machine learning predictions, including GWAS, across several simulated genomic scenarios mimicking homoplasic, oligogenic and polygenic traits with varying effect sizes and heritability, using as test organism *K. pneumoniae*. Although we mostly tested our framework on simulated quantitative traits, we also applied it to real MICs of four representative antibiotics Ciprofloxacin, Gentamicin, Piperacillin-Tazobactam and Meropenem.

The results on simulated data showed that for classification the overall performance decreases as the number of classes increases, as expected due to the increased difficulty of exact classification. For this reason, we also tested the performance of the models by 1-tier accuracy, allowing one-either-side predicted class to be considered as correct. The Random Forest method resulted in more reliable handling predictions in the vast majority of the simulations (1-tier accuracy range 0.86-1), highlighting this algorithm as more suitable when dealing with class imbalance. We also addressed how the application of censoring decreases the accuracy of the models, suggesting how it should be avoided as it prevents them from having the total amount of information available. Dealing with the regression model, the Elastic Net and Random Forest achieved overall similar results over different levels of *h^2^*, increasing the performances when the MIC were not binned but treated as numerical values.

Concerning the prediction of real MICs, including estimation of heritability reported for the first time in *K. pneumoniae*, we observed how both the heritability and performances of the models varied across different antibiotics, showing a better fit of the classification model.

Additionally, to evaluate the reliability of the model for detection of causal variants set in the simulations we performed multiple GWAS analysis, showing Elastic Net, Random Forest and FaST-LMM as able to detect the causative variant used in the homoplastic simulation, albeit the number of false negatives was highly influenced by the tuning of the parameters. Whether multiple causative loci are involved, the Random Forest exhibited a more conservative behaviour reducing the total false positive compared to Elastic Net and FaST-LMM, prioritising only variants with higher effects.

We therefore recommend that a practical solution to handle MICs data is to consider each case separately, evaluating the genetic architecture of each resistance trait (which causative variants are already known), the number of antibiotic-step concentrations available, and accounting for the balance of the MIC bins when they are treated as categories. When a small number of antibiotic-step concentrations is available (3-7), considering the MICs as categorical is preferrable, and is also valuable to binarize prediction in case of few concentrations. Alternatively, as the number of classes increases, the regression-based approach represents a valid option as well as off-by-one errors in classification accuracy, depending on what false positive/false negative trade-off is desired.

Our work presents some limitations. Firstly, we only relied on genes and SNPs, and did not consider additional more general genetic features such as unitigs, which may improve the performance of the models, as they provide more genomic information than SNPs [28, 49]. Moreover, using SNPs can result in a higher number of false negatives when the causal variants are not located in core positions but in plasmids, suggesting that SNPs may not be ideal for gene based resistance prediction. Moreover, there can be a lack of single SNPs able to discriminate between adjacent MICs concentrations [24]. While our simulations covered a broad range of scenarios, we only used a single real dataset focusing only on one species and four antibiotics. Furthermore, we did not consider prior information (e.g. distribution of effect sizes of variants / previously known relations with resistance traits) in our model [24, 28, 30], as possible implementation within a Bayesian framework via an ordinal regression model.

In conclusion, collections of high-quality genomes are increasingly populating global databases, and availability of MIC data would be equally recommended, possibly specifying details of the phenotypic test used. Subsequently, laboratory tests able to increase the range of antibiotic-step concentrations (e.g. E-test) should be considered — reducing the level of censored data — when building these models. Indeed, while it is easier to indicate a binary phenotype, this interpretation is not consistent over time, and results in permanent information loss. Having more information-rich phenotype data would allow more modelling possibilities, and especially as datasets grow may help improve prediction accuracy in future, as the number of samples increases far beyond what we have been able to study here.

## Author contributions

Methodology: G.B.B, L.C., J.A.L. Conceptualization: G.B.B, J.A.L. and D.S. Formal analysis: G.B.B., J.L. Writing-original draft: G.B.B. and J.A.L. Writing – review and editing: G.B.B., J.A.L., D.S., M.C., E.J.F., L.C.

