## Supplementary material for "Optimising machine learning prediction of minimum inhibitory concentrations in *Klebsiella pneumoniae*"

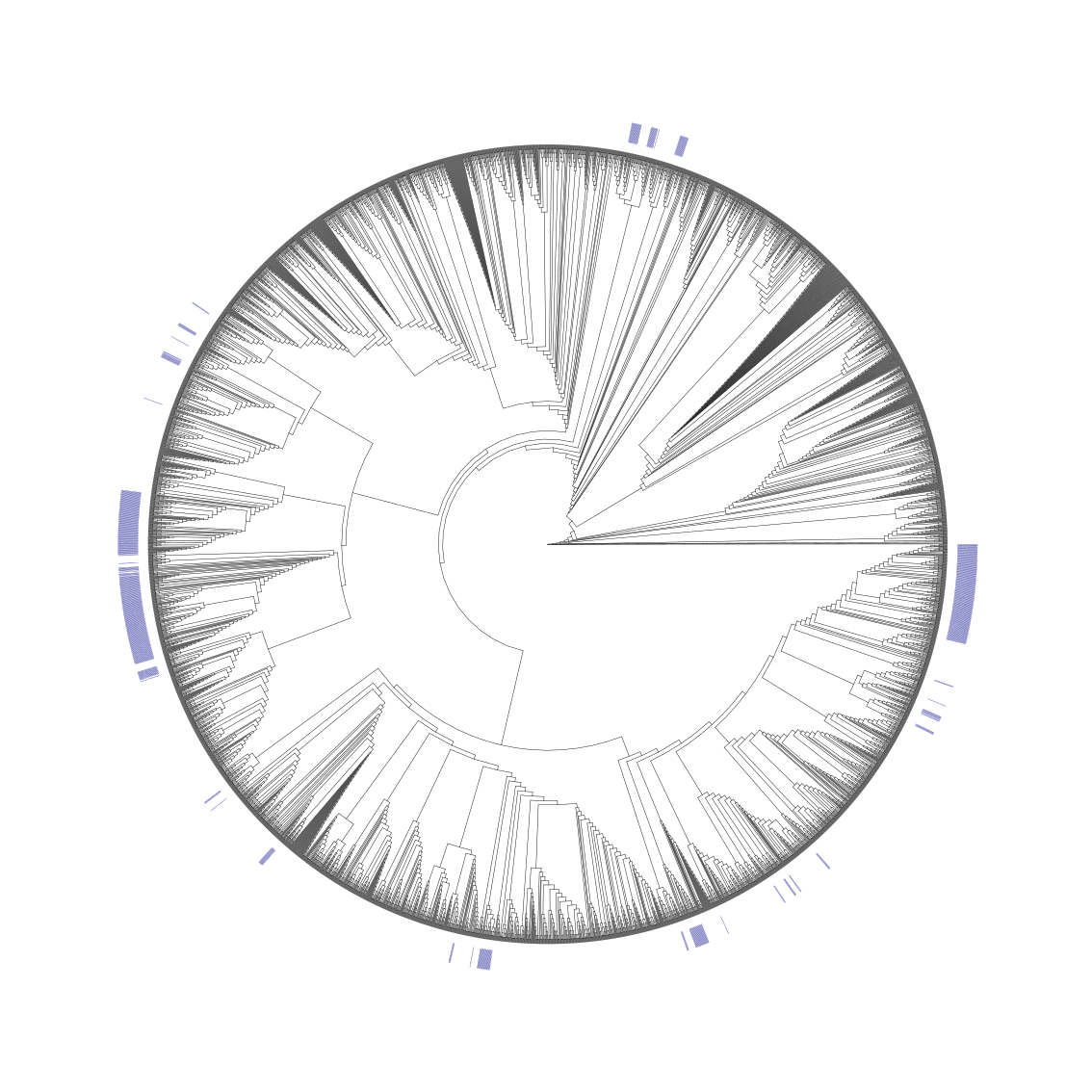

Supplementary Figure 1) Phylogenetic tree representing the distribution of the homoplastic SNP in the *maoA* gene chosen for the simulation.

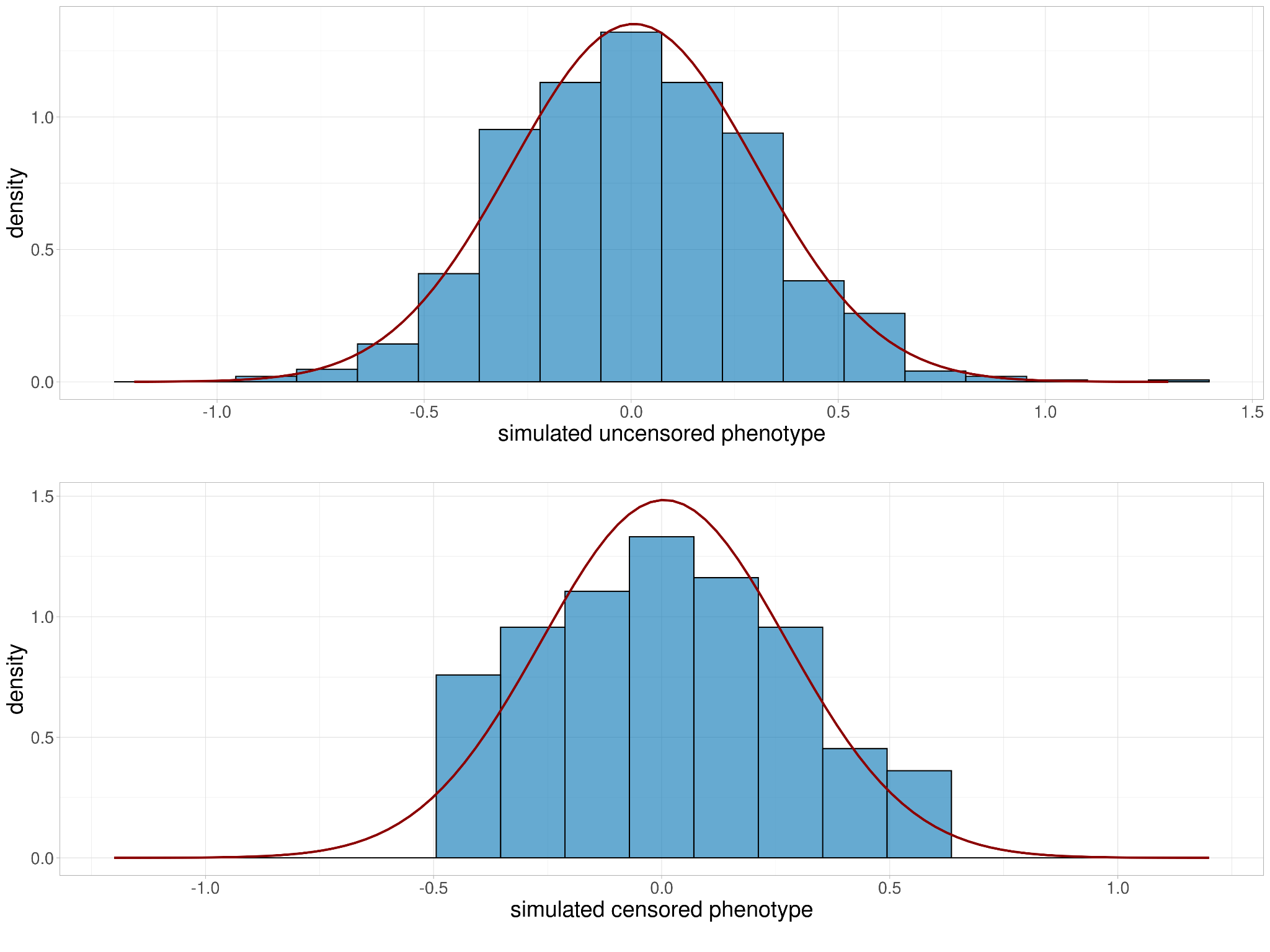

Supplementary Figure 2) General representation of normal distribution before and after applying censoring to resemble MIC measurement in reality. The censoring is applied based on quantile intervals.

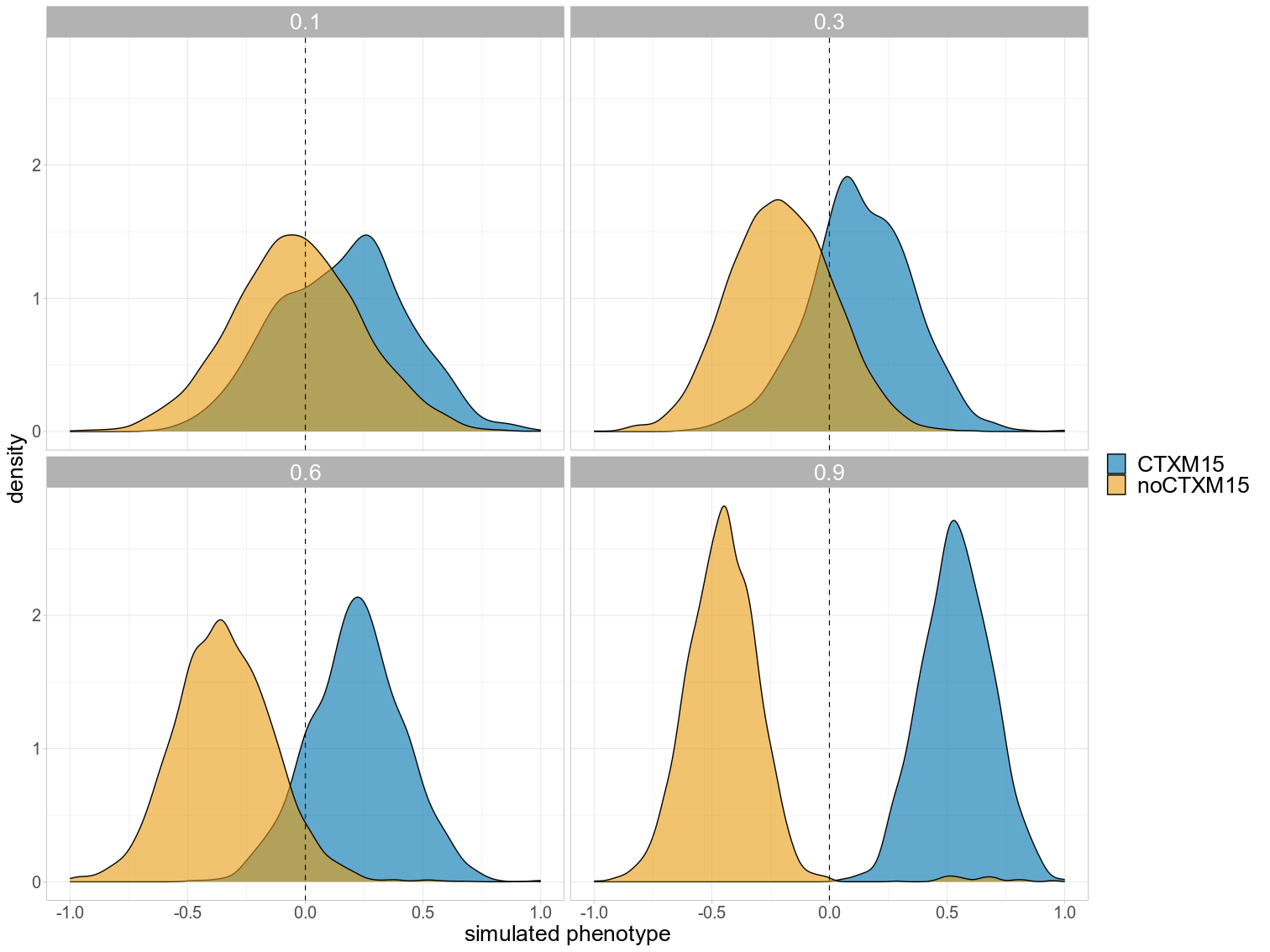

Supplementary Figure 3) Simulated phenotype distribution using homoplastic genes as causal variants at different levels of *h²*. The areas are coloured according to the presence-absence of the causal variant.

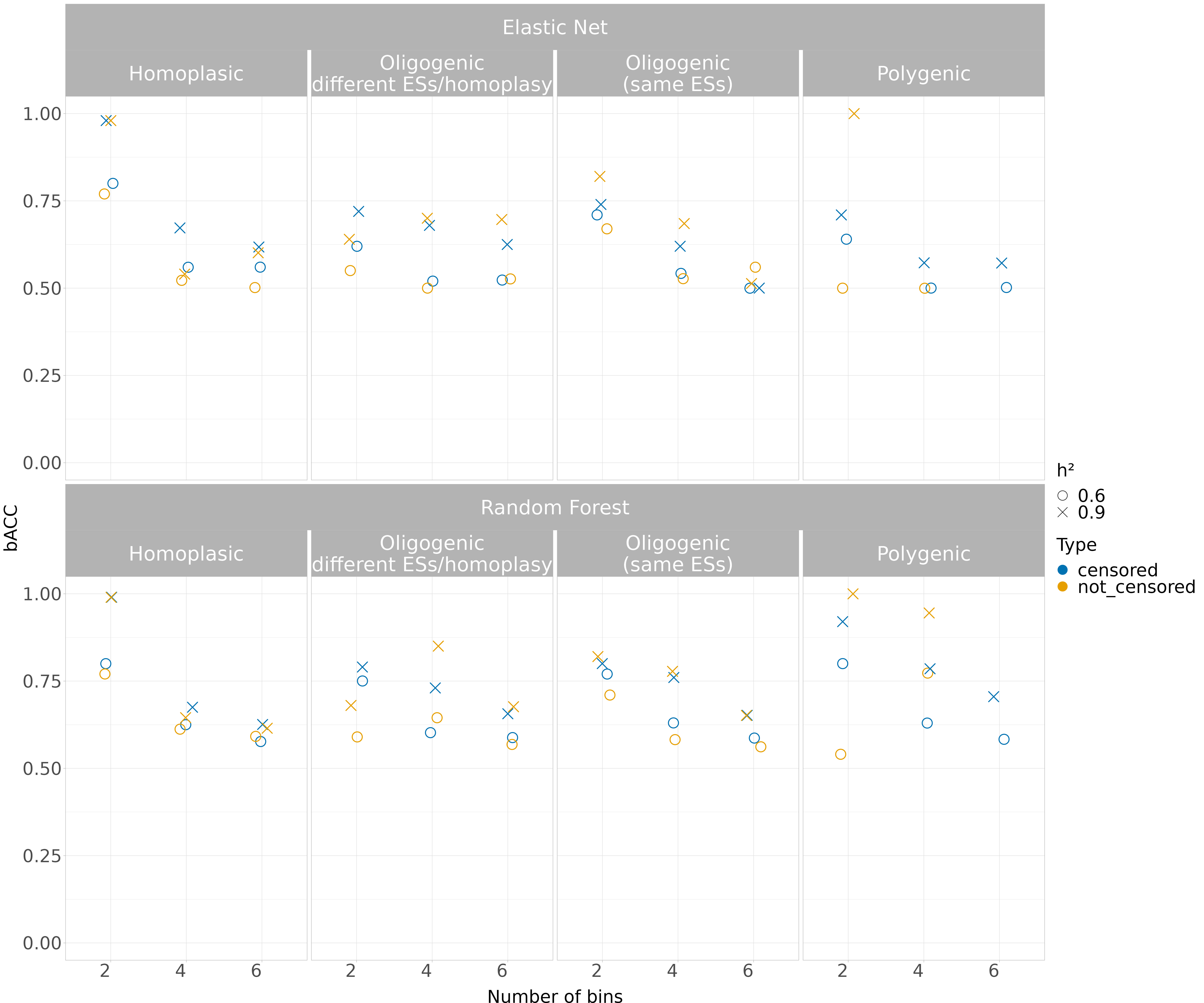

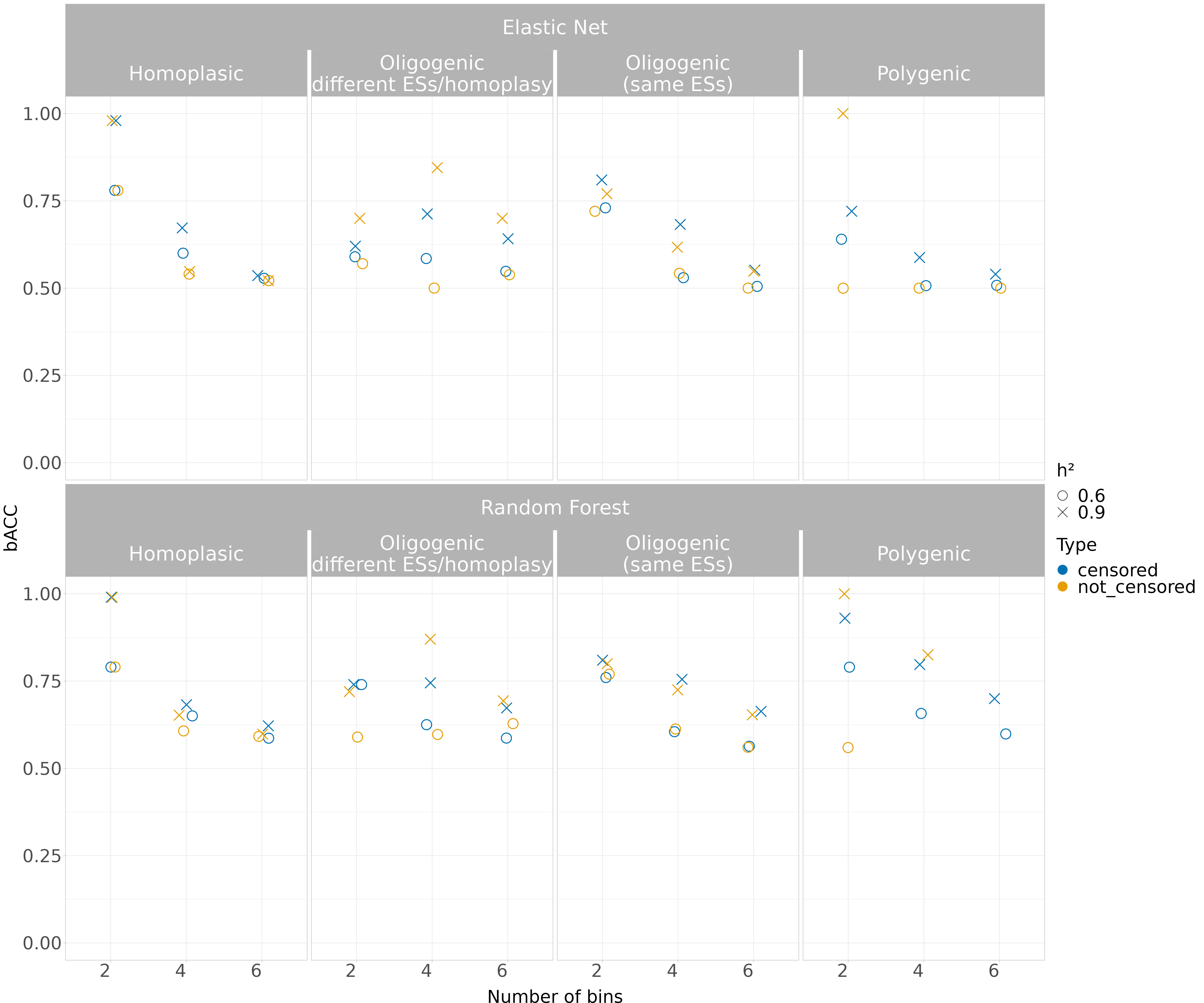

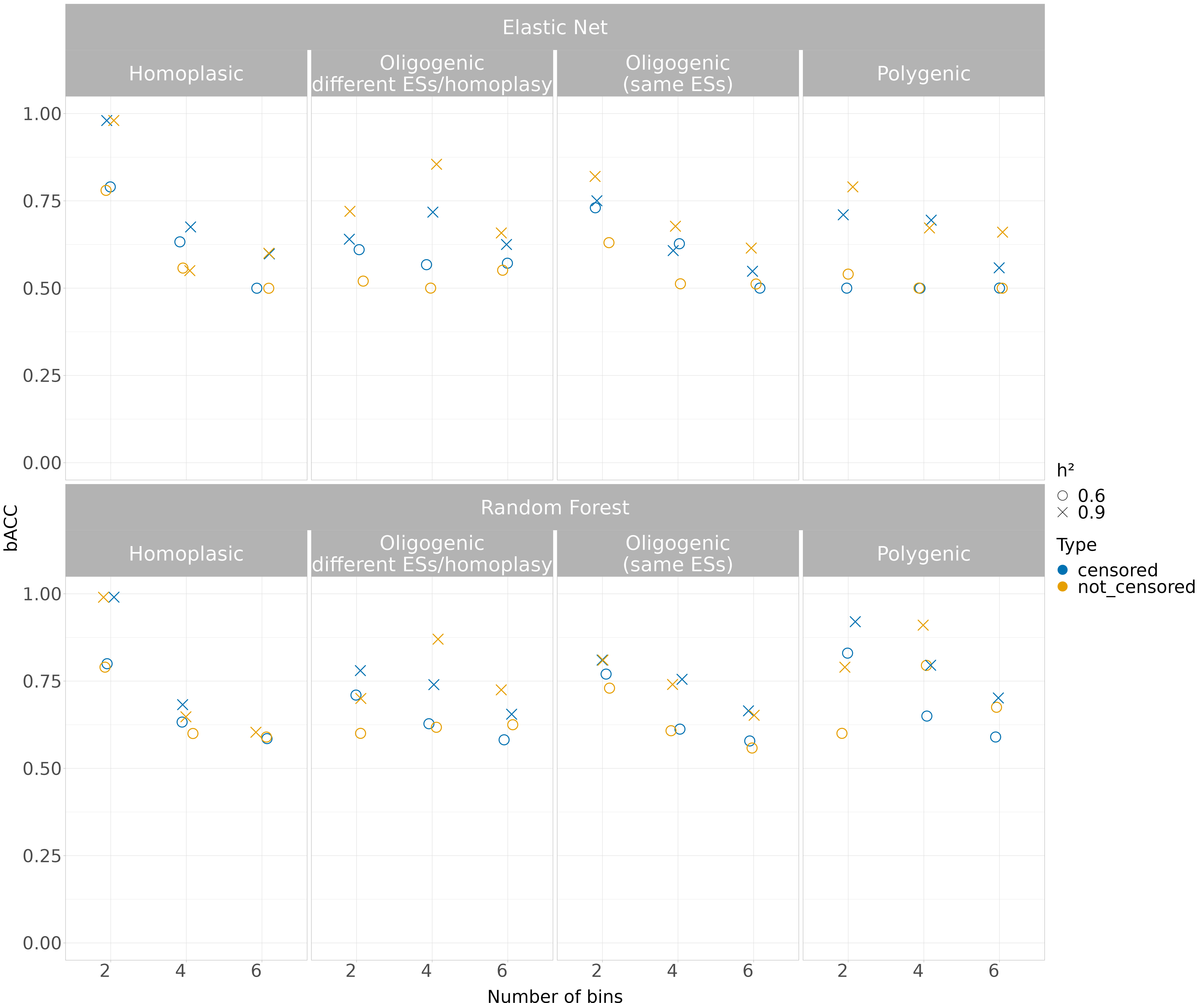

Supplementary Figure 4) Performance of the classification models (effect size = 1.5 except for the oligogenic simulation) measured using balanced accuracy (bACC), indicating the arithmetic mean of sensitivity and specificity. The models were benchmarked over two levels of *h²*, considering both the censored and not censored binned simulated traits.

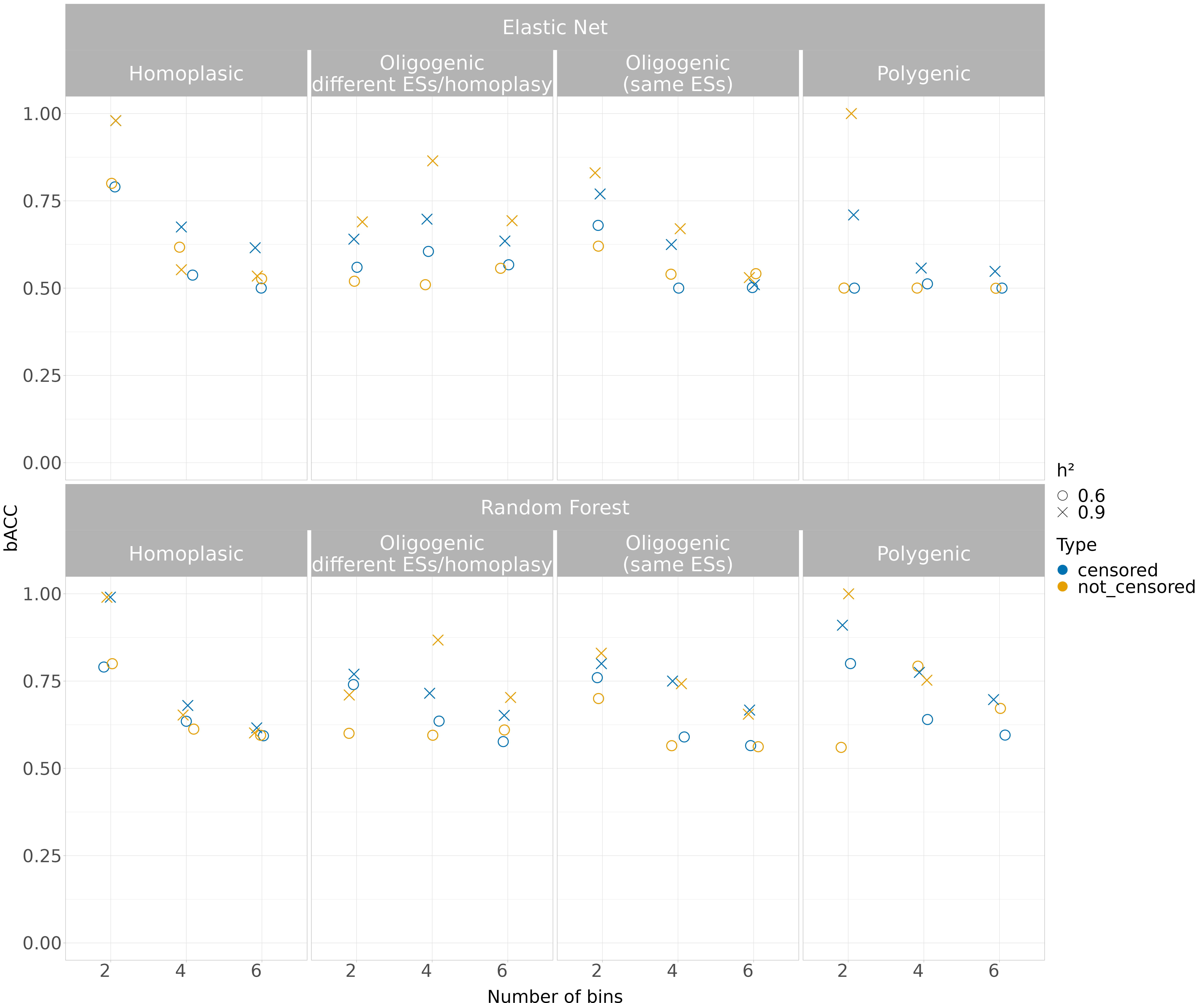

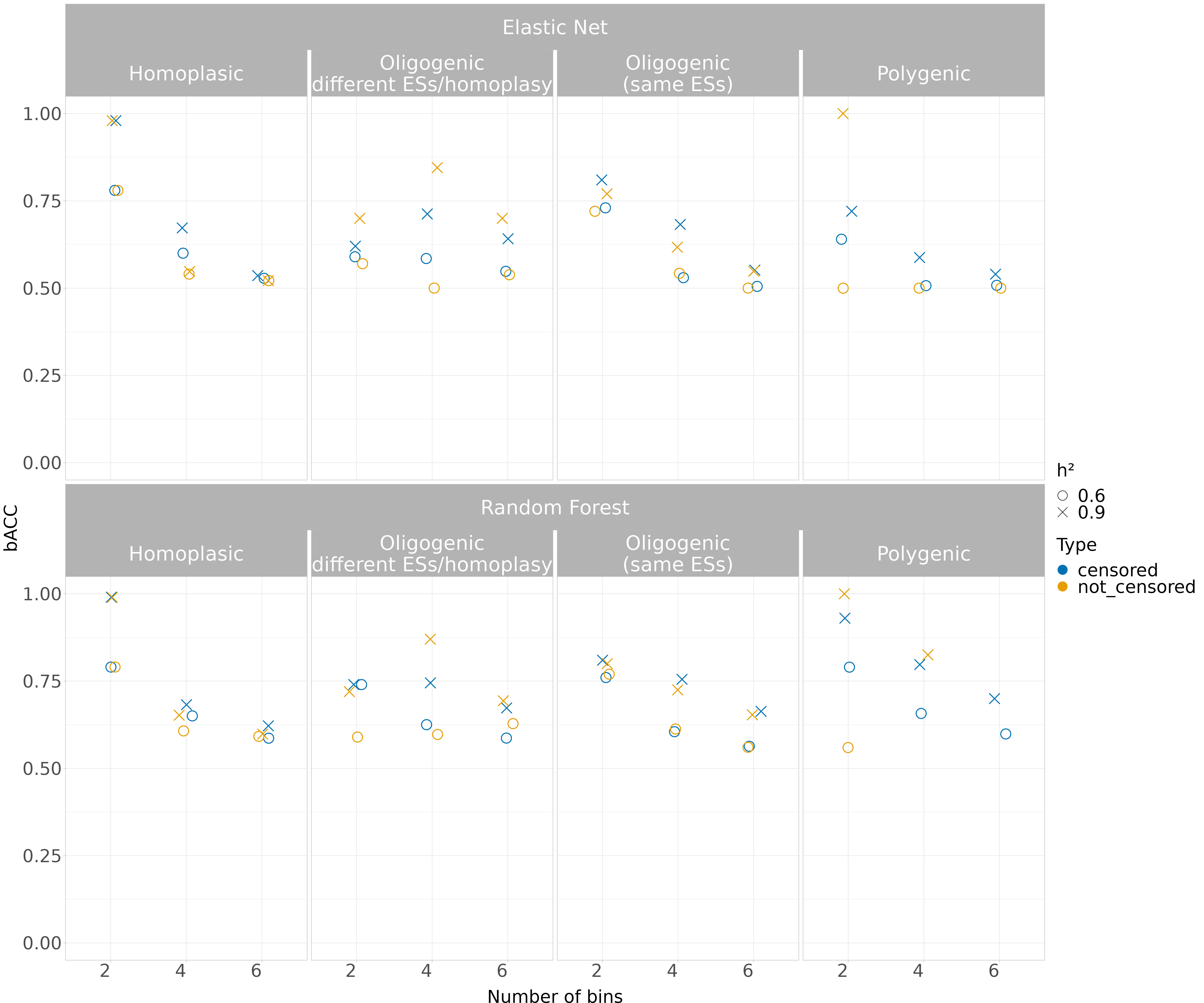

Supplementary Figure 5) Performance of the classification models (effect size = 10 except for the oligogenic simulation) measured using balanced accuracy (bACC), indicating the arithmetic mean of sensitivity and specificity. The models were benchmarked over two levels of *h²*, considering both the censored and not censored binned simulated traits.

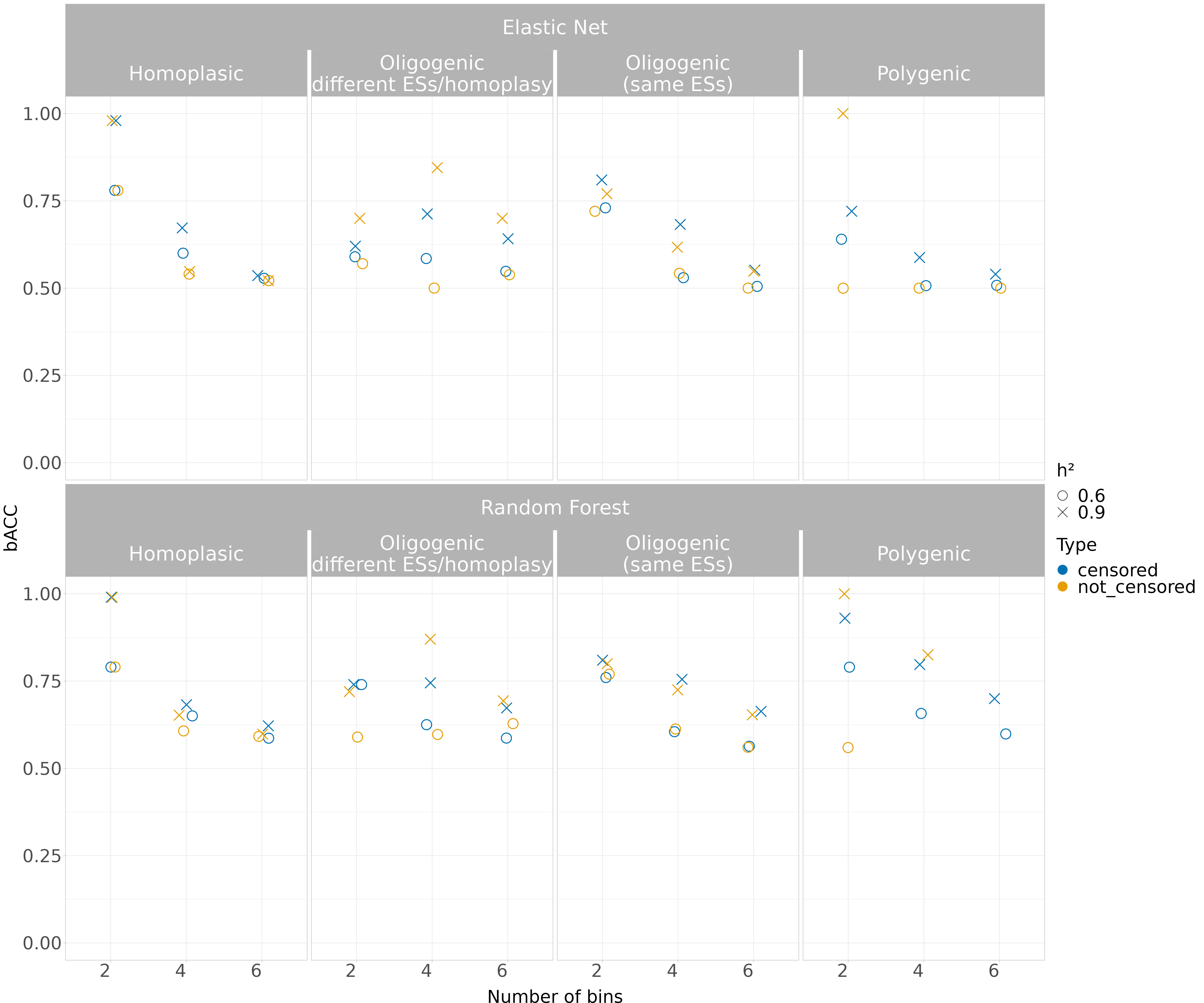

Supplementary Figure 6) Performance of the classification models (effect size = 30 except for the oligogenic simulation) measured using balanced accuracy (bACC), indicating the arithmetic mean of sensitivity and specificity. The models were benchmarked over two levels of *h²*, considering both the censored and not censored binned simulated traits.

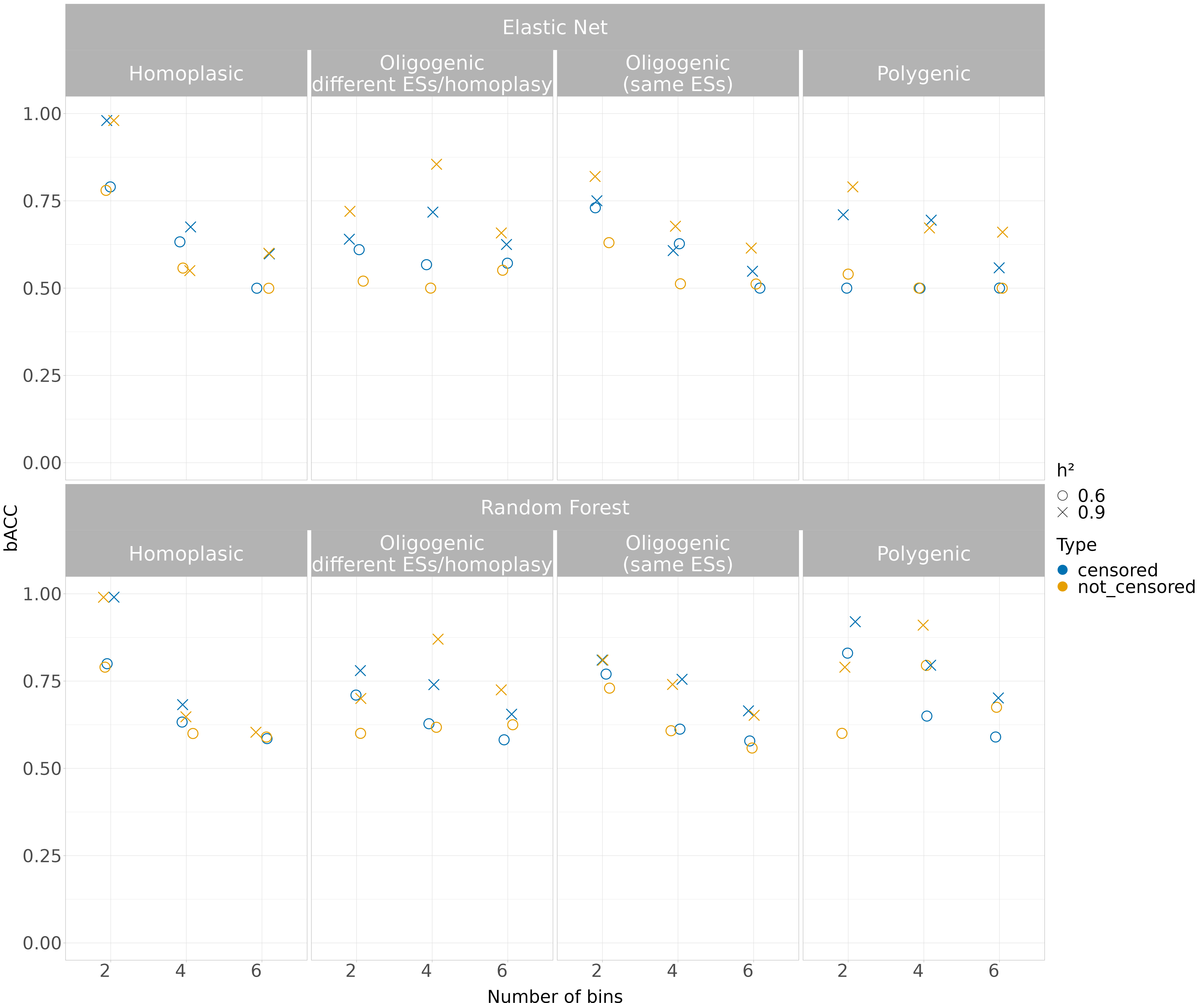
Supplementary Figure 7) Performance of the classification models (effect size = 100 except for the oligogenic simulation) measured using balanced accuracy (bACC), indicating the arithmetic mean of sensitivity and specificity. The models were benchmarked over two levels of *h²*, considering both the censored and not censored binned simulated traits.

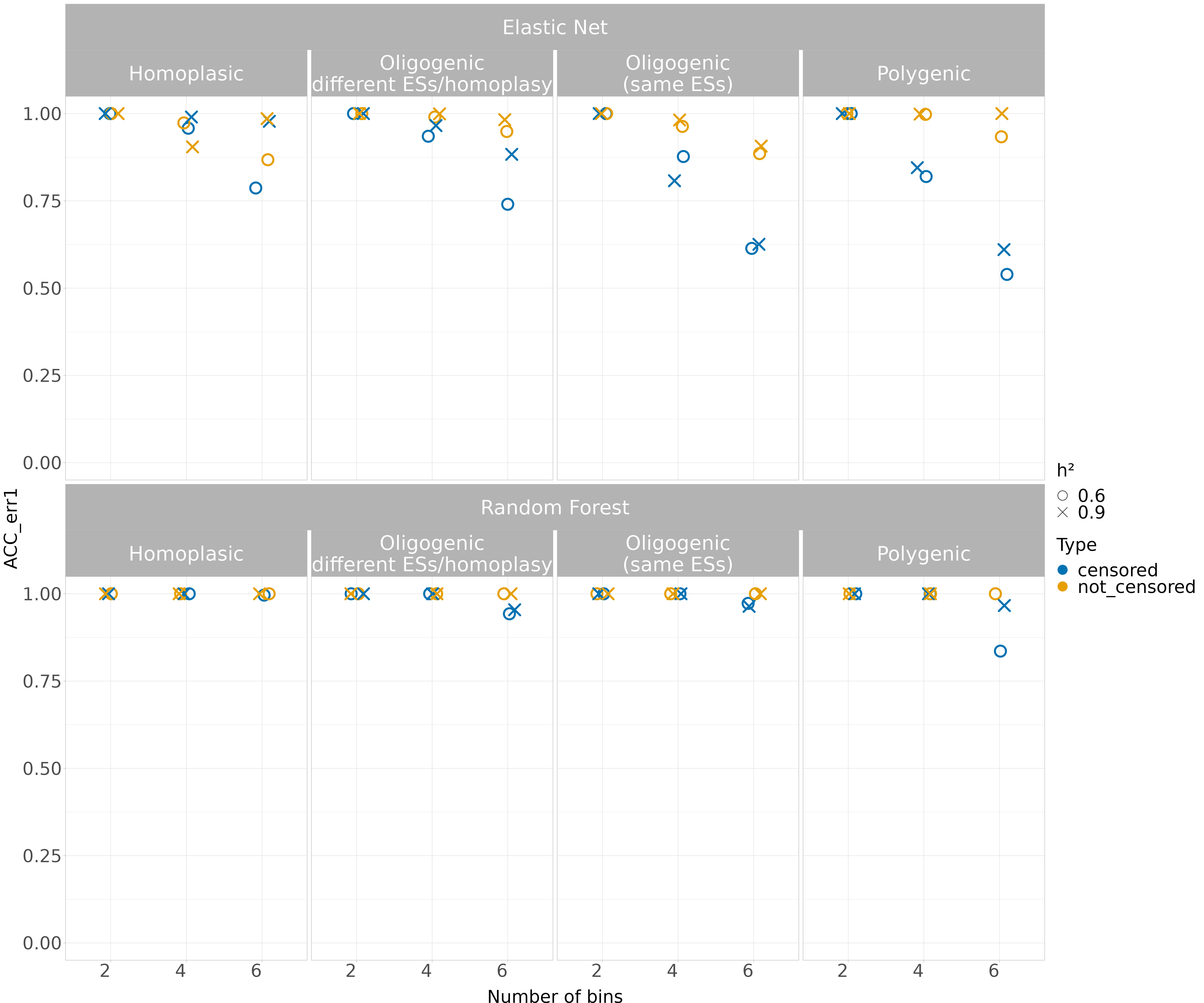
Supplementary Figure 8) Performance of the classification models (effect size = 1.5 except for the oligogenic simulation) when the accuracy (need to correct y-axis label) was adjusted to consider one-either-side predicted class as correct (accuracy within ± 1 two-fold dilution factor). The models were benchmarked over two levels of *h²*, considering both the censored and not censored binned simulated traits.

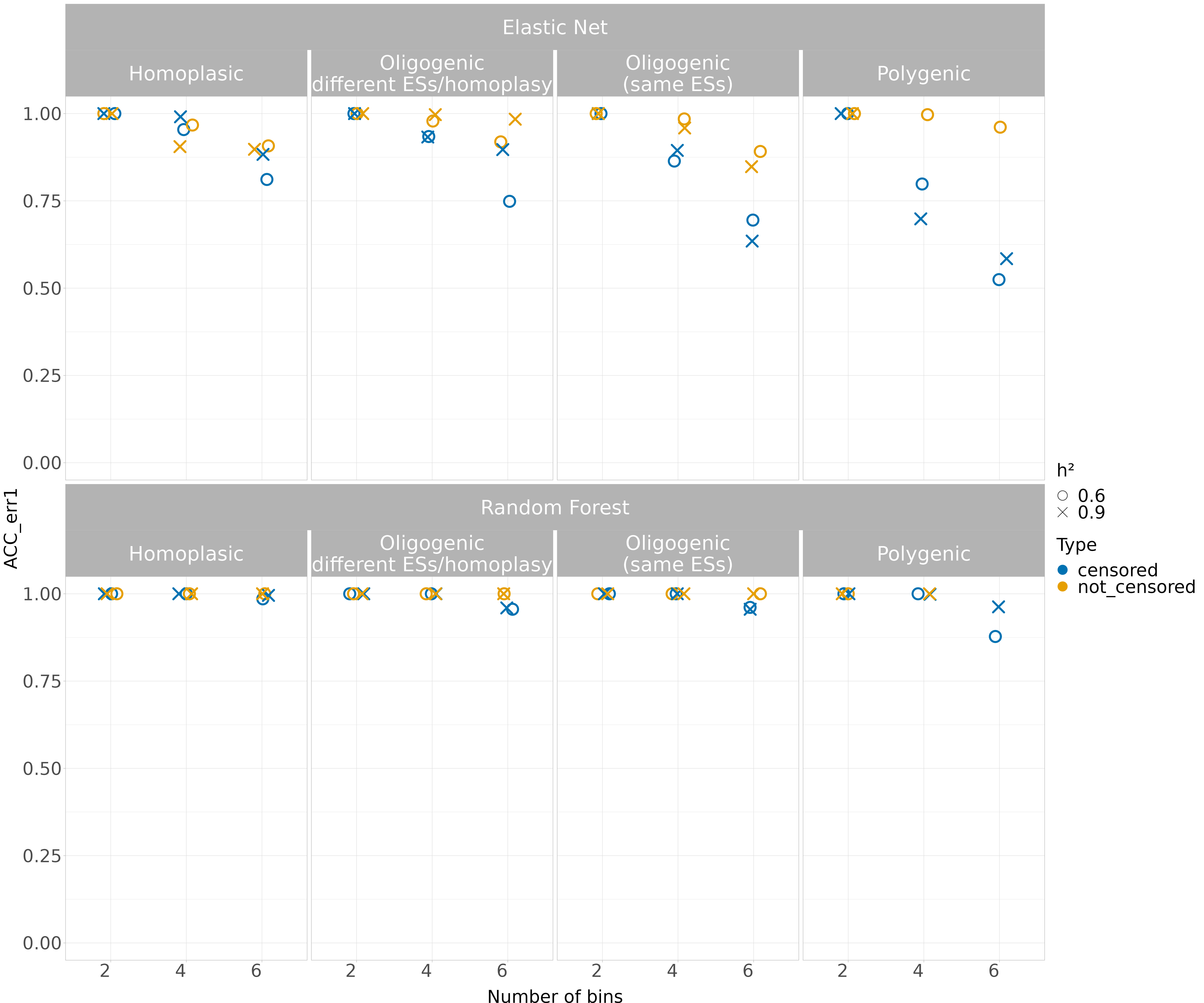
Supplementary Figure 9) Performance of the classification models (effect size = 10 except for the oligogenic simulation) when the accuracy (need to correct y-axis label) was adjusted to consider one-either-side predicted class as correct (accuracy within ± 1 two-fold dilution factor). The models were benchmarked over two levels of *h²*, considering both the censored and not censored binned simulated traits.

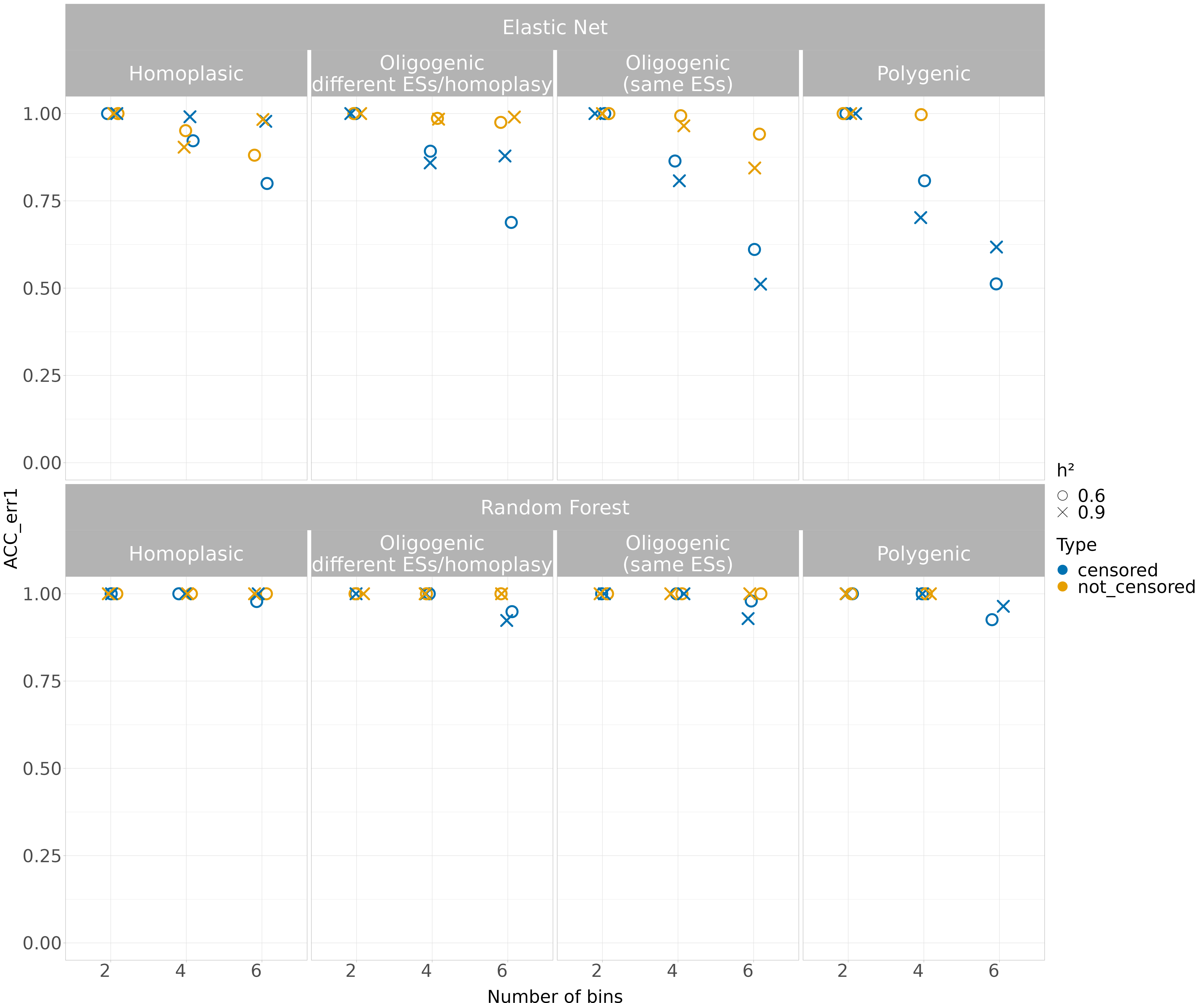

Supplementary Figure 10) Performance of the classification models (effect size = 30 except for the oligogenic simulation) when the accuracy (need to correct y-axis label) was adjusted to consider one-either-side predicted class as correct (accuracy within ± 1 two-fold dilution factor). The models were benchmarked over two levels of *h²*, considering both the censored and not censored binned simulated traits.

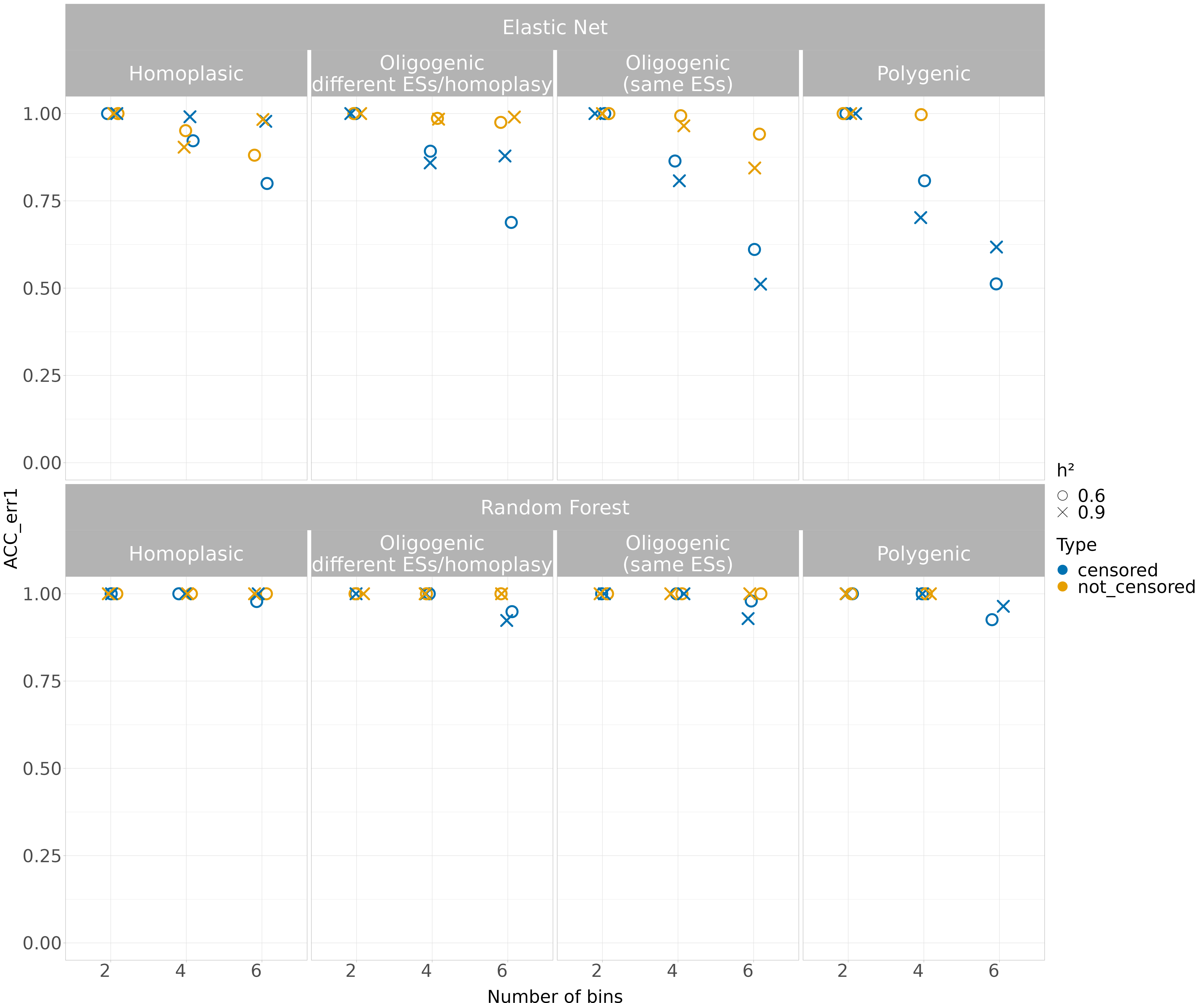
Supplementary Figure 11) Performance of the classification models (effect size = 100 except for the oligogenic simulation) when the accuracy (need to correct y-axis label) was adjusted to consider one-either-side predicted class as correct (accuracy within ± 1 two-fold dilution factor). The models were benchmarked over two levels of *h²*, considering both the censored and not censored binned simulated traits.

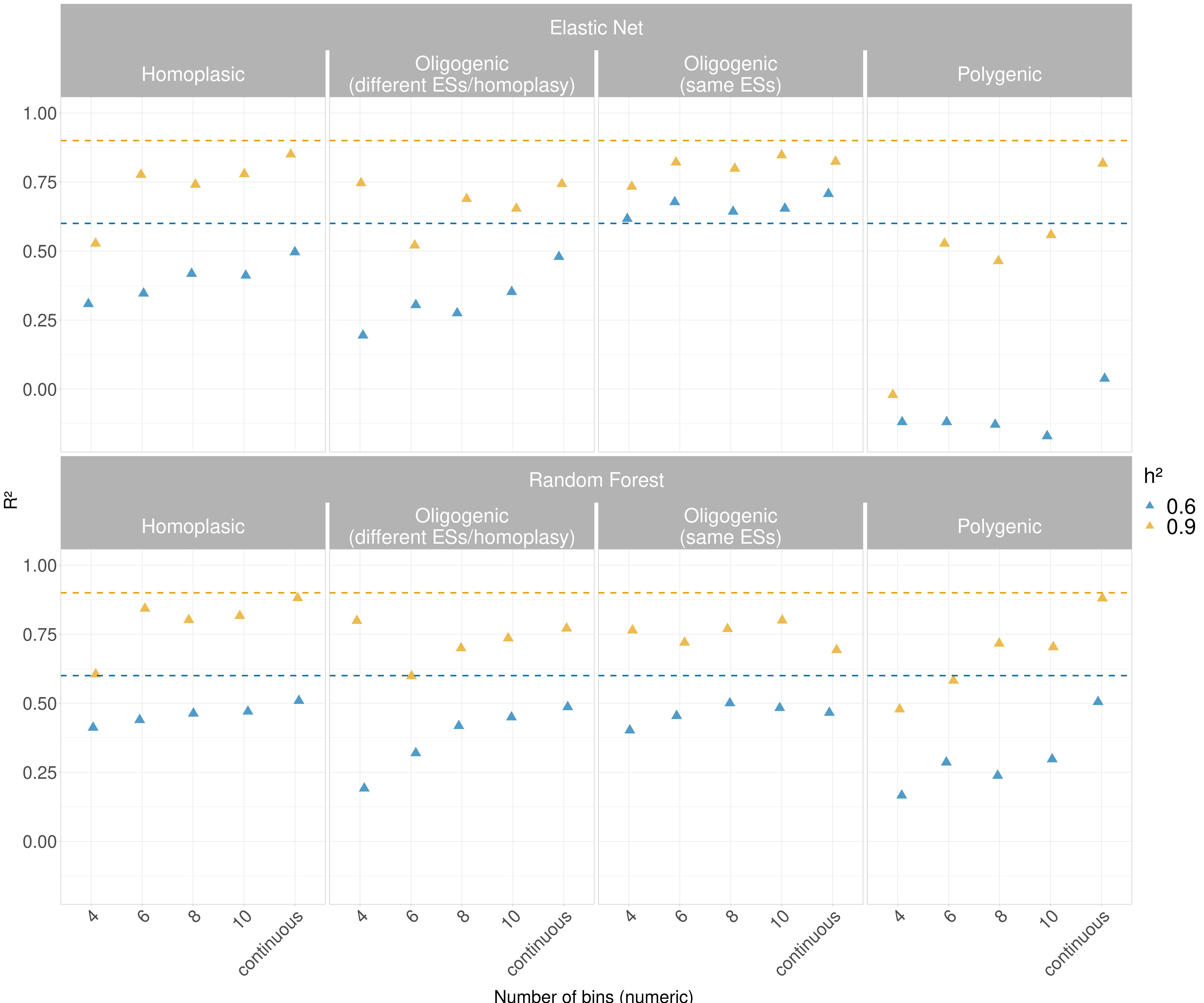
Supplementary Figure 12) Performance of regression model (effect size = 1.5 except for the oligogenic simulation) between Elastic Net and Random Forest when dealing with simulated MICs. The bins obtained from the simulation were binned into multiple intervals and treated as numeric. Also, the simulated quantitative traits without binning were used. Since the *h²* is the proportion of phenotypic variance explained by genotype, and thus an equivalent of the *R²* in regression, the dashed lines are used to assess the capability of the models to estimate the *h²*.

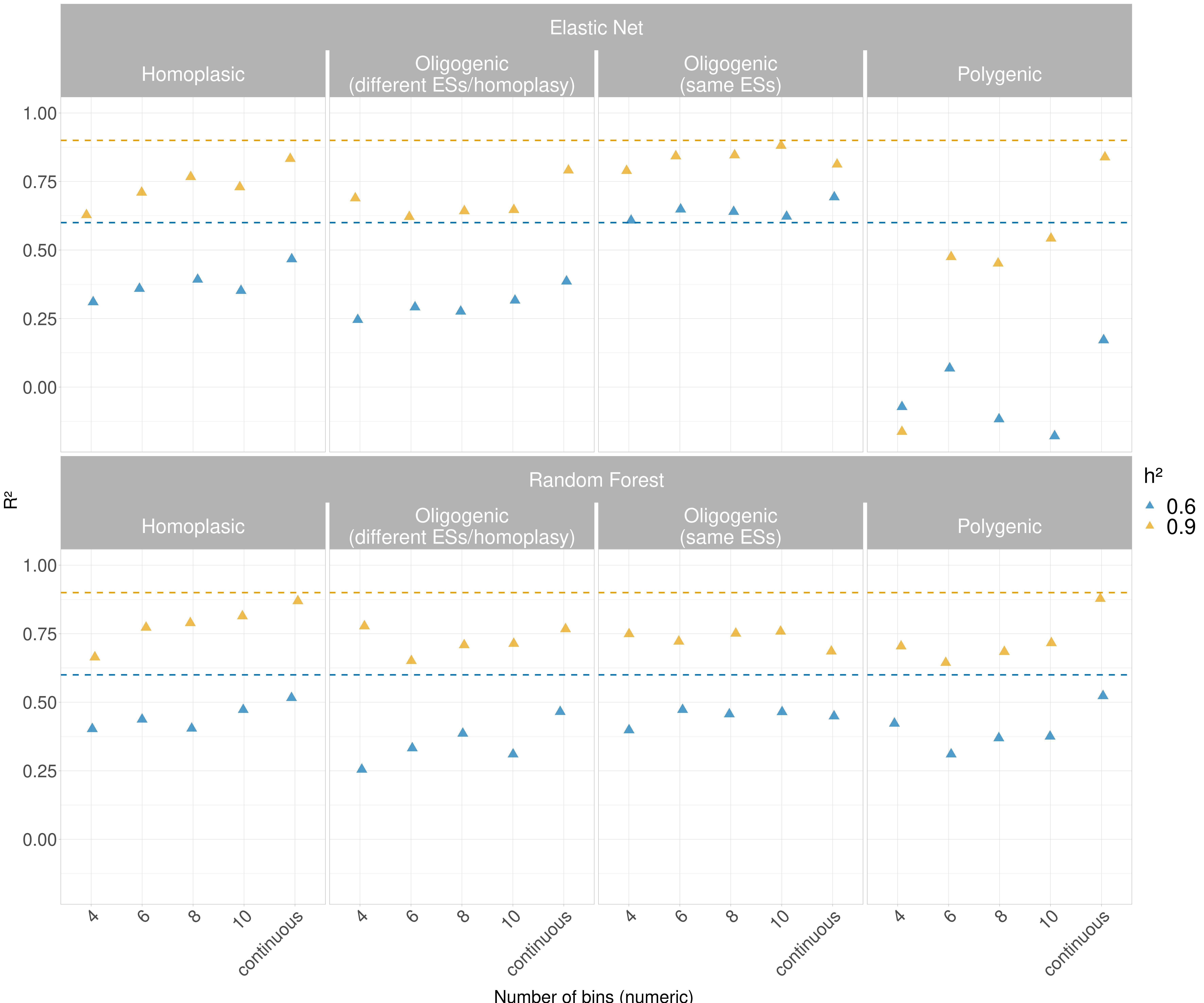
Supplementary Figure 13) Performance of regression model (effect size = 10 except for the oligogenic simulation) between Elastic Net and Random Forest when dealing with simulated MICs. The bins obtained from the simulation were binned into multiple intervals and treated as numeric. Also, the simulated quantitative traits without binning were used. Since the *h²* is the proportion of phenotypic variance explained by genotype, and thus an equivalent of the *R²* in regression, the dashed lines are used to assess the capability of the models to estimate the *h²*.

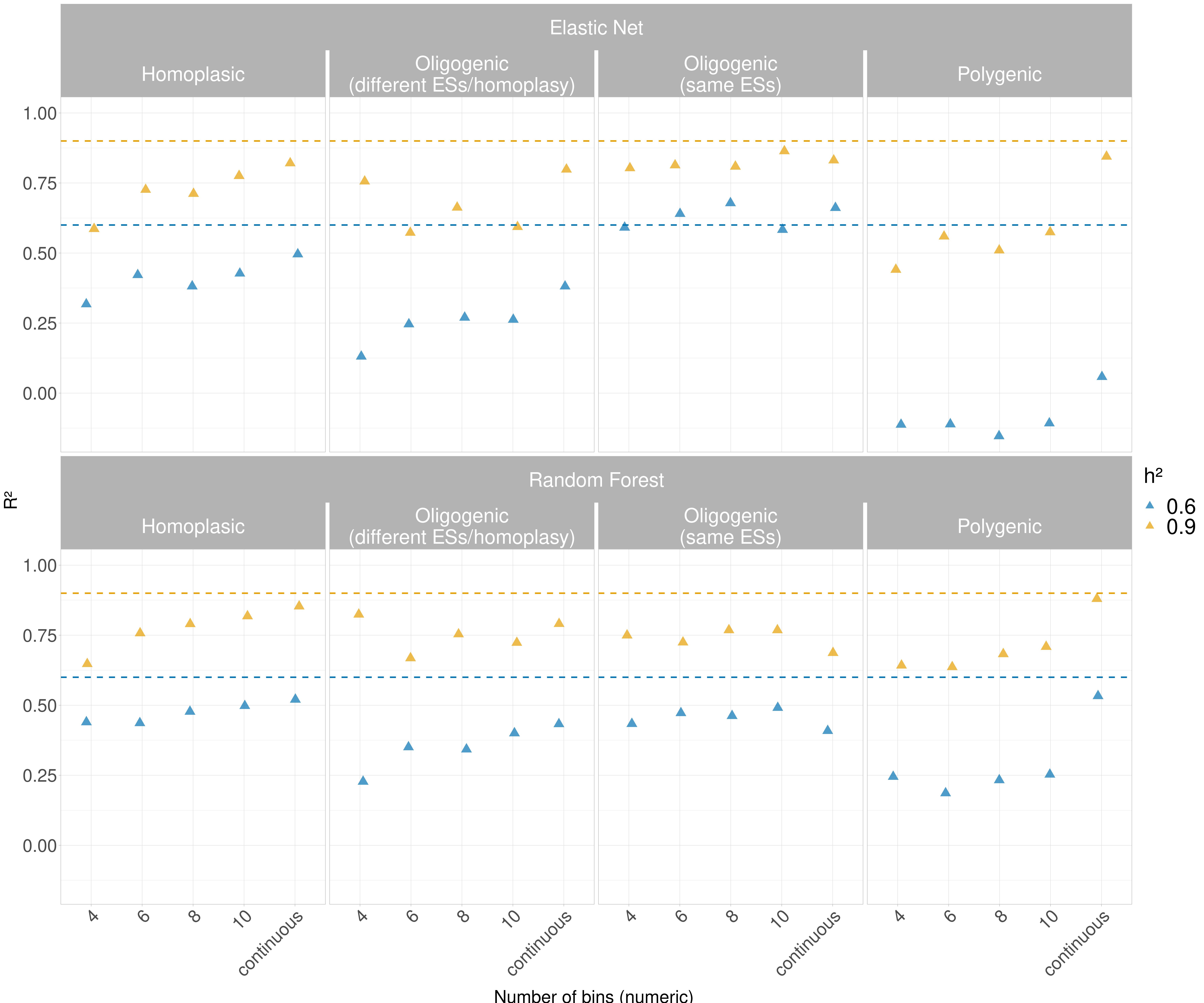
Supplementary Figure 14) Performance of regression model (effect size = 30 except for the oligogenic simulation) between Elastic Net and Random Forest when dealing with simulated MICs. The bins obtained from the simulation were binned into multiple intervals and treated as numeric. Also, the simulated quantitative traits without binning were used. Since the *h²* is the proportion of phenotypic variance explained by genotype, and thus an equivalent of the *R²* in regression, the dashed lines are used to assess the capability of the models to estimate the *h²*.

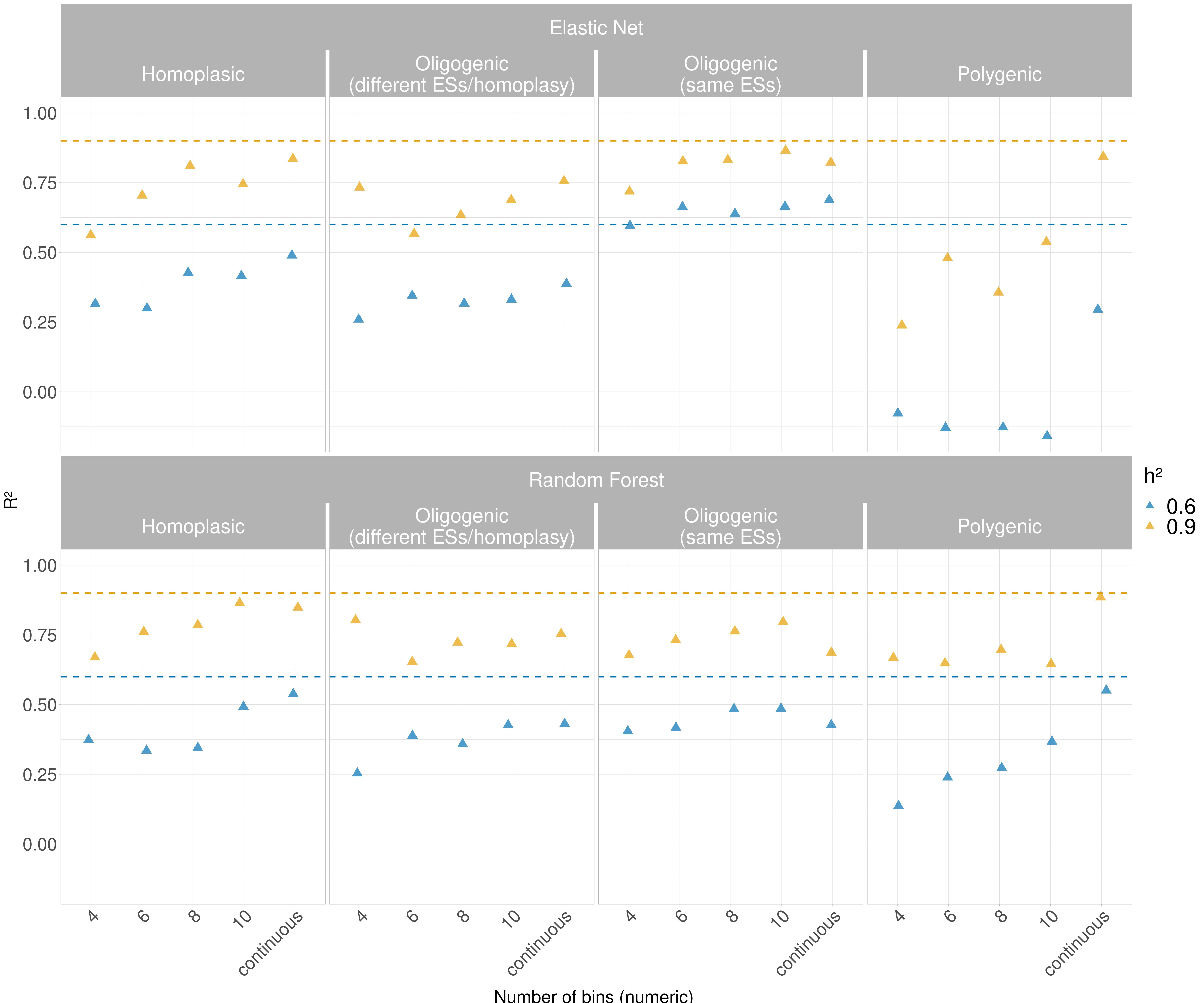

Supplementary Figure 15) Performance of regression model (effect size = 100 except for the oligogenic simulation) between Elastic Net and Random Forest when dealing with simulated MICs. The bins obtained from the simulation were binned into multiple intervals and treated as numeric. Also, the simulated quantitative traits without binning were used. Since the *h²* is the proportion of phenotypic variance explained by genotype, and thus an equivalent of the *R²* in regression, the dashed lines are used to assess the capability of the models to estimate the *h²*.

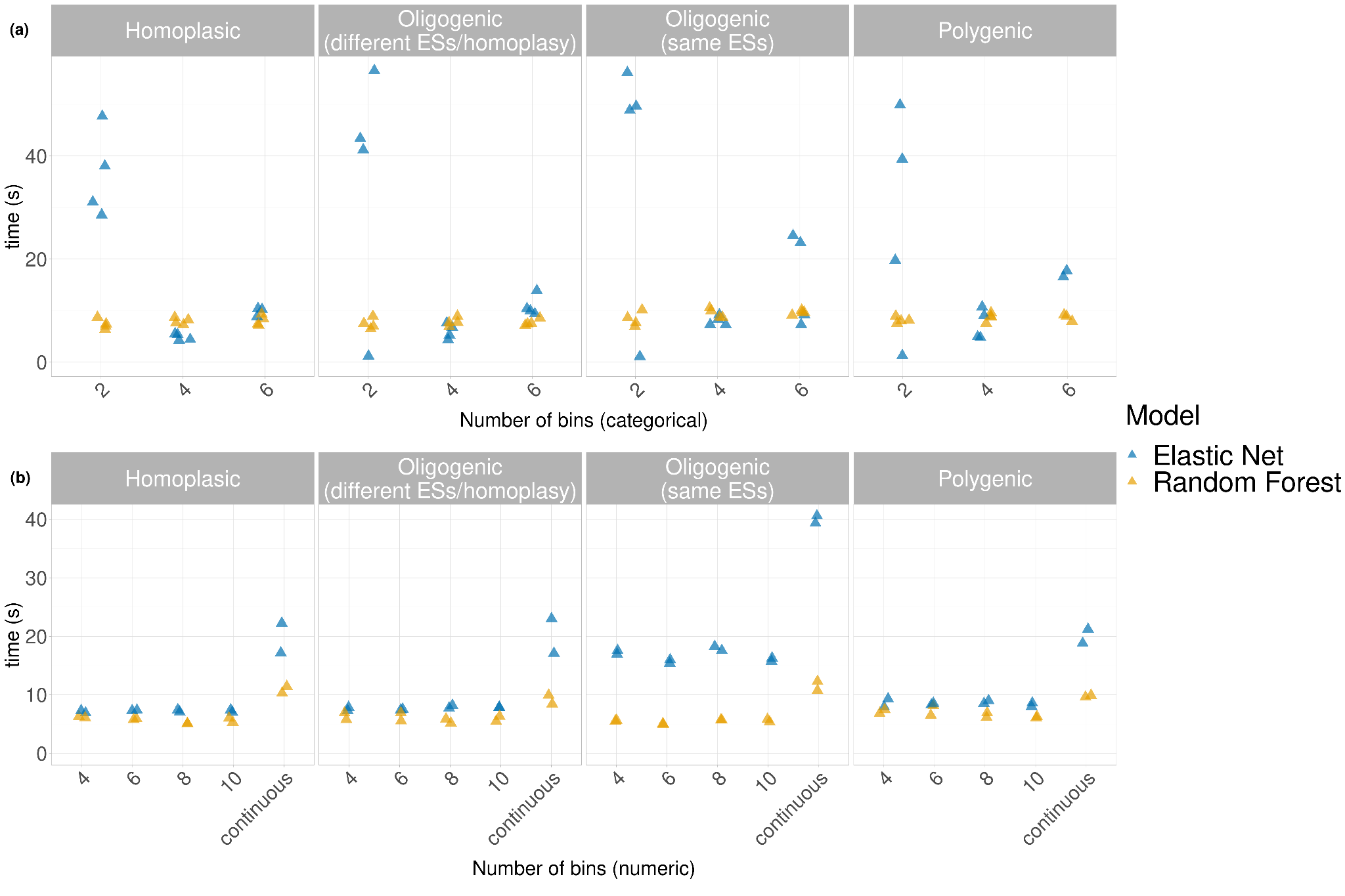

Supplementary Figure 16) Time required for training and prediction (seconds) of the two models for a) classification and b) regression.

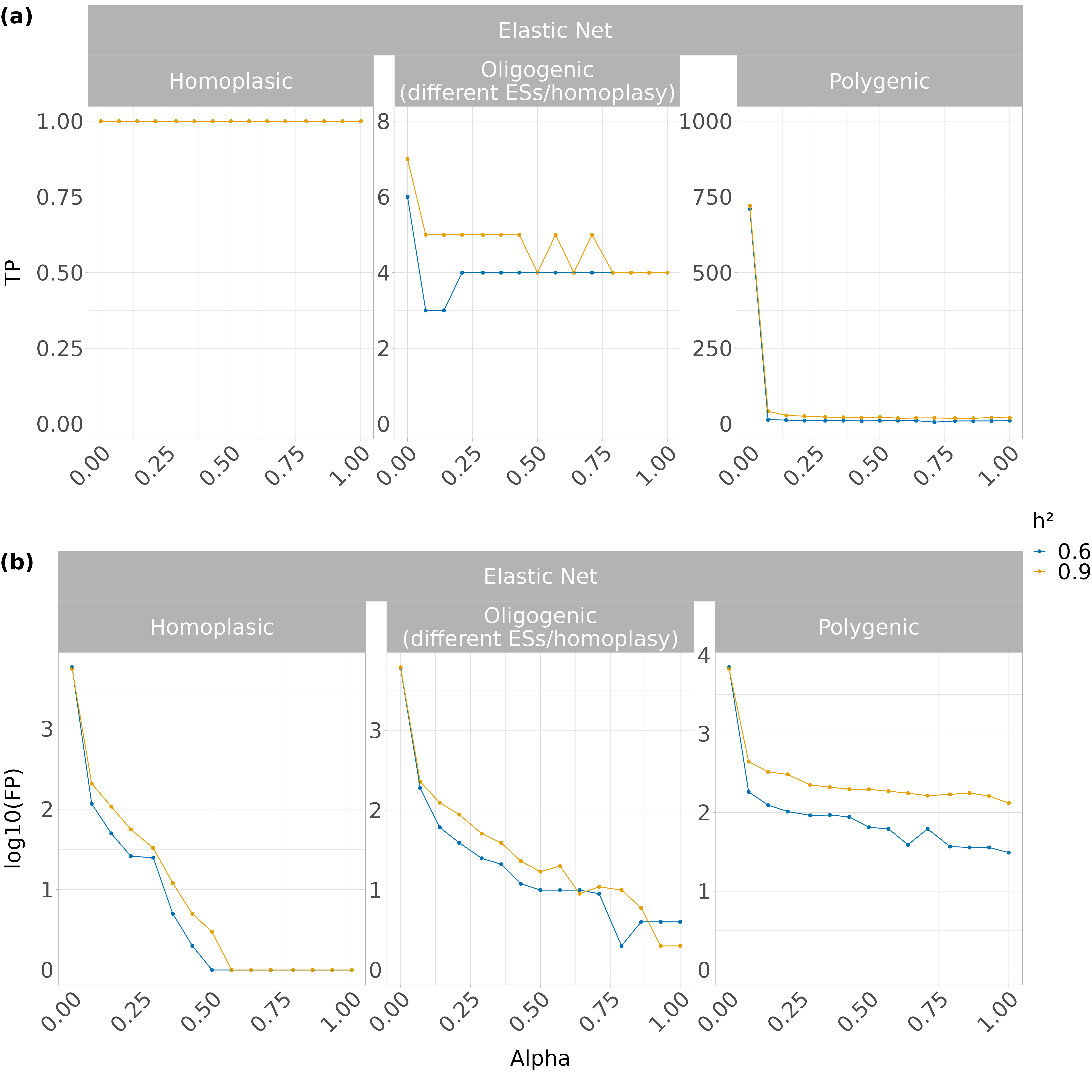

Supplementary Figure 17) Elastic Net accuracy to detect causal variants. The variation of true and false positives was assessed through the tuning of the α value, ranging from α=0 (Ridge) to α=1 (LASSO) a) Total number of true positives possible (i.e. causal variants): 1 (homoplasic); 8 (oligogenic); 1000 (polygenic). b) Total number of false positives possible: 12410 (homoplasic); 12403 (oligogenic); 11403 (polygenic).

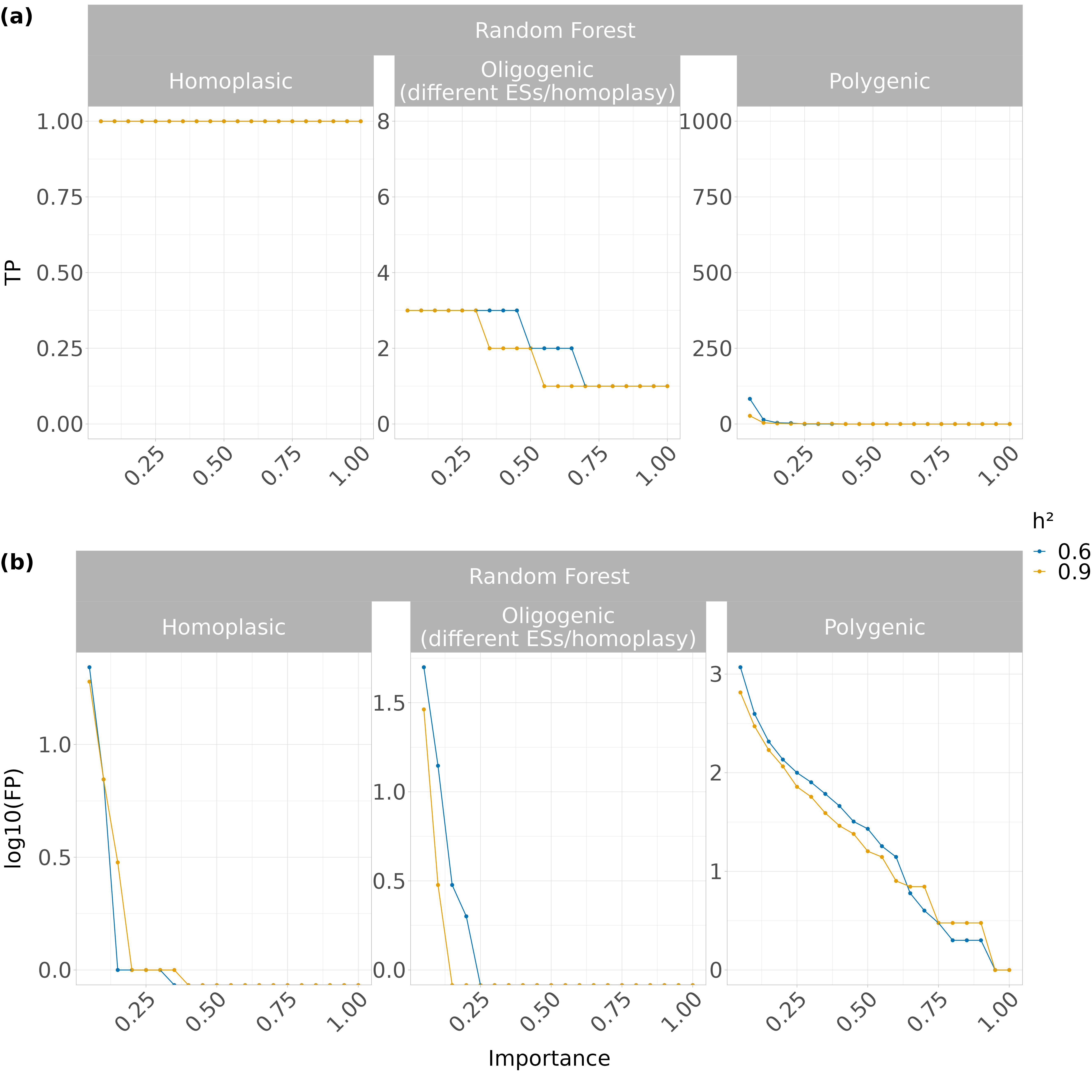
Supplementary Figure 18) Random Forest accuracy to detect causal variants. The feature importance parameter was normalised between 0-1 and used to detect the variation of true and false positives depending on the cut-off. a) Total number of true positives possible (i.e. causal variants): 1 (homoplasic); 8 (oligogenic); 1000 (polygenic). b) Total number of false positives possible: 12410 (homoplasic); 12403 (oligogenic); 11403 (polygenic).

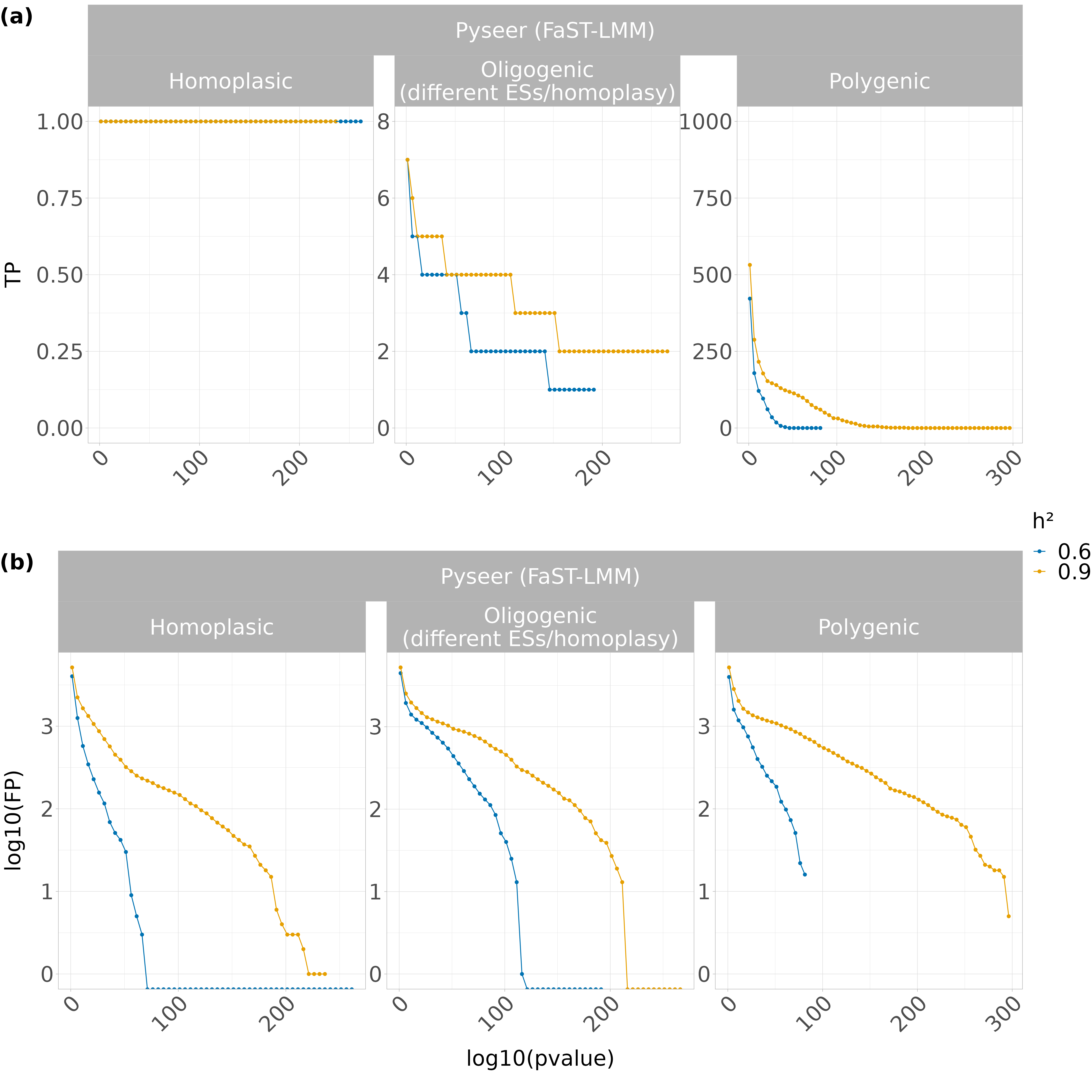
Supplementary Figure 19) FaST-LMM (Pyseer) accuracy to detect causal variants. The log10 (p-value) was used to detect the number of true and false positives depending on the selected cut-off. a) Total number of true positives possible (i.e. causal variants): 1 (homoplasic); 8 (oligogenic); 1000 (polygenic). b) Total number of false positives possible: 12410 (homoplasic); 12403 (oligogenic); 11403 (polygenic).
